# Multigenerational Proteolytic Inactivation of Restriction Upon Subtle Genomic Hypomethylation

**DOI:** 10.1101/2024.10.21.619520

**Authors:** Esther Shmidov, Alexis Villani, Senén D. Mendoza, Ellay Avihu, Ilana Lebenthal-Loinger, Sarit Karako-Lampert, Sivan Shoshani, Chang Ye, Yiding Wang, Hao Yan, Weixin Tang, Joseph Bondy-Denomy, Ehud Banin

## Abstract

Restriction-modification (R-M) systems, present in most bacterial genomes, protect against phage infection by detecting and degrading invading foreign DNA. However, like many prokaryotic anti-phage systems, R-M systems pose a significant risk of auto-immunity, exacerbated by the presence of hundreds to thousands of potential cleavage sites in the bacterial genome. In *Pseudomonas aeruginosa*, restriction inactivation upon growth at high temperatures was previously described, however, which system is being inactivated, the underlying mechanism, as well as the timing of recovery, remain unknown. Here, we report that *P. aeruginosa* Type I methyltransferase (HsdMS) and restriction endonuclease (HsdR) components are degraded by two Lon-like proteases when replicating above 41 °C, which induces partial genome hypomethylation and simultaneously prevents self-targeting, respectively. Interestingly, upon return to 37 °C, methyltransferase activity returns gradually, with restriction activity not fully recovering for over 60 bacterial generations, representing the longest bacterial memory to our knowledge. Forced expression of HsdR over the first 45 generations is toxic, demonstrating the fitness benefit of HsdR inactivation. Our findings demonstrate that type I R-M is tightly regulated post-translationally with a remarkable memory effect to ensure genomic stability and emphasize the importance of mitigating auto-toxicity for bacterial defense systems.

## Introduction

Bacteriophages (phages) are viruses that infect bacteria. Over millions of years of co-evolution, bacterial strains have evolved various anti-phage defense systems^1^. The first discovered genes to have anti-phage activity were the Restriction-Modification (R-M) systems^2^. However, despite their early discovery, many fundamental aspects of R-M system biology remain poorly understood. To date, over 100 additional defense systems have been identified^3^, varying in the mechanism of action, phage targets, and viral sensing^4^. Alongside their protective role, bacterial defense systems often feature nucleases or other potentially toxic enzymes, generating a risk for autoimmunity^5^ and need to be tightly regulated to become activated only for defense purposes.

R-M systems protect the bacterial cell from phage infection by detecting and degrading invading foreign DNA. Most bacterial genomes (83%) carry at least one R-M system^6^. These systems modify the bacterial genome at system-specific modification sites with the methyltransferase component and subsequently degrade the viral genome with the restriction endonuclease holoenzyme upon detection of unmodified sites in the invading DNA. In terms of autoimmunity and toxicity, R-M systems hold a great risk as the restriction motifs are located within the bacterial chromosome in high numbers. One example of autoimmunity avoidance from type I R-M systems in *Escherichia coli* is restriction alleviation (RA), where the protease ClpXP degraded the endonuclease to prevent self-targeting after DNA damage^7,8,9^. For *Pseudomonas aeruginosa*, an opportunistic gram-negative pathogen^10^, type I R-M represents one of the most abundant defense systems among the isolates^11^. In the PAO1 laboratory strain, a commonly used laboratory strain of P. aeruginosa, an active type I R-M system is encoded by the three proximal genes PA2735 (*hsdM)*, PA2734 (*hsdS*), and PA2732 (*hsdR*) with a modification pattern of the GATC(N)_6_GTC sequence^12,13^. HsdMS form the active methyltransferase while the HsdMSR holoenzyme is the endonuclease^14^ (Fig. 1A).

**Figure 1:**
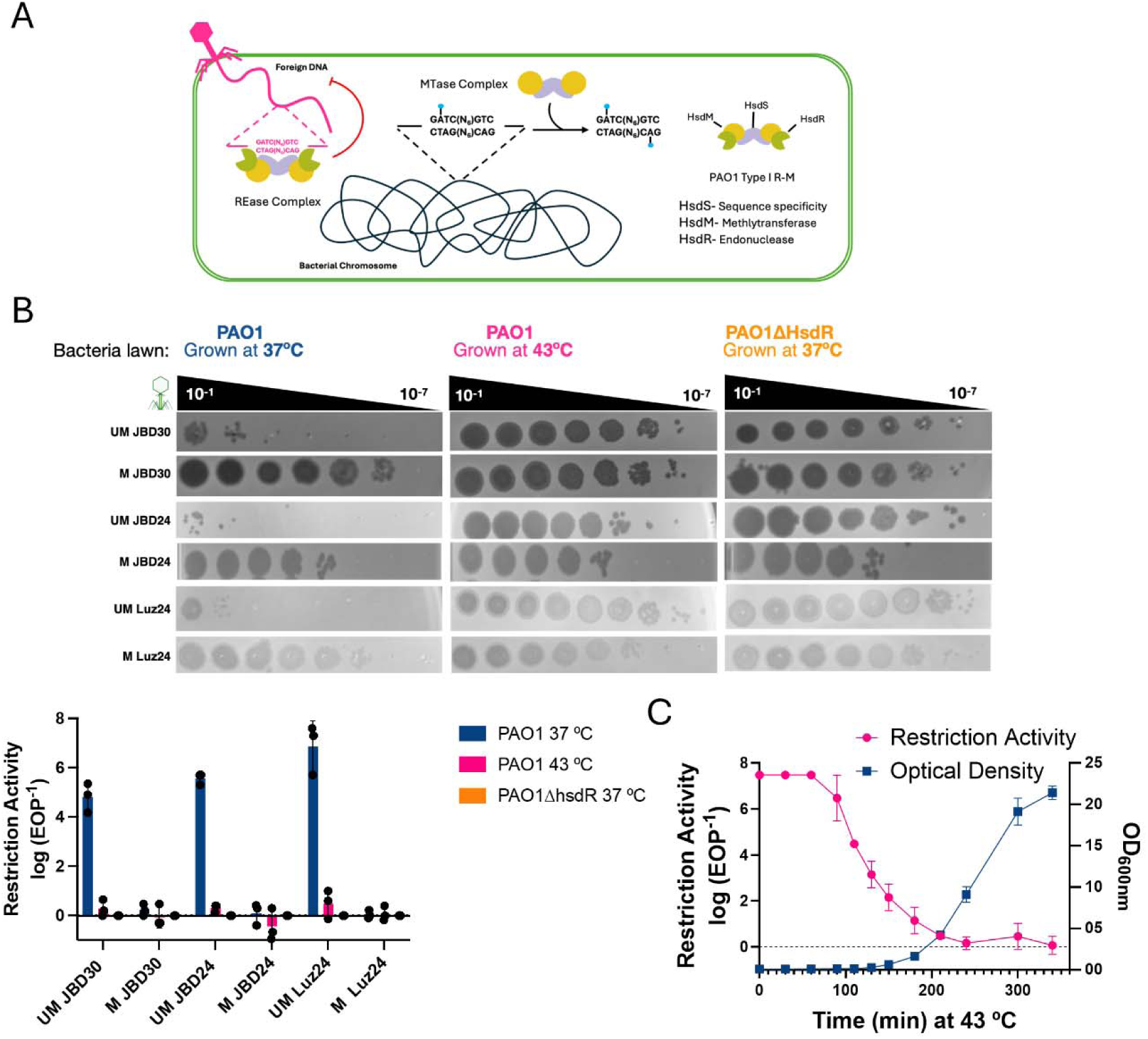
Elevated growth temperature leads to inactivation of restriction endonuclease. **(A)** Schematic representation of Type I R-M system. **(B)** Comparative plaquing efficiency and restriction activity of WT grown overnight at either 37 °C or 43 °C and restriction mutant (Δ*hsdR*) at 37 °C infected with either UM or M phages. **(C)** Log-phase and replication are required for iREN: Restriction activity was examined for PAO1 WT strain during growth at 43 °C at intervals parallel to growth curve measurements through optical density (OD 600nm). The graphs in (B) and (C) represent the average of three independent biological replicates, with error bars indicating standard deviation.

In 1965, Holloway reported the inactivation of restriction activity when *P. aeruginosa* was grown at 43 °C^15^. However, which R-M system was being controlled in this experiment, and the mechanism of action are not understood. Here, we show that the Type I R-M is inactivated in *P. aeruginosa* upon multiple rounds of cell division and nascent DNA synthesis at elevated temperatures (>41 °C), which induces genomic hypomethylation. Notably, transient heat shock, DNA damage, or prolonged exposure to high temperatures during the stationary phase do not induce restriction inactivation. By studying the effects of culturing *P. aeruginosa* at elevated temperatures as a model for the consequences of hypomethylation, we demonstrate that methylation deficiency is indeed necessary to induce restriction loss and that Lon proteases are responsible for a partial decrease in the levels of the methyltransferase HsdMS and a strong inactivation of HsdR. Upon return to lower temperatures (i.e. 37 °C), restriction remarkably remains inactive for up to 60 generations and, for most of this period, forced expression of HsdR is toxic. We named this phenomenon iREN for **i**nactivation of **R**estriction **En**donuclease.

## Results

### Bacterial replication at 43 °C leads to complete inhibition of the type I R-M system

We have previously shown that the Type I R-M system in strain PAO1 is active against phage JBD30^13^. Therefore, we first set out to establish that the previously observed loss of restriction activity in 1965^15^ in a strain described as “PAO” could be recapitulated and connected to this specific system. To establish this connection, we used JBD30 (containing 5 restriction sites), a related temperate phage JBD24 (6 sites), and the distinct lytic phage Luz24 (5 sites). Modified phages (M) were propagated in the WT PAO1 strain, while unmodified phages (UM) were generated by propagating them in strain PA14, which lacks any Type I R-M system. When UM phages were used to infect WT PAO1 (grown at 37 °C), their efficiencies of plating (EOP) were reduced by approximately 5 orders of magnitude, compared to a Δ*hsdR* strain, indicating successful restriction (Fig. 1B). To examine the inactivation of restriction activity, we grew overnight (ON) the PAO1 WT strain, at 43 °C instead of 37 °C. Cells grown at 43 °C showed complete inactivation of restriction <u>en</u>donuclease (which we refer to as the iREN state), phenocopying the Δ*hsdR* mutant. These results demonstrate that the Type I R-M system in PAO1 is inactivated during growth at high temperature.

We next investigated the requirements for the establishment of iREN. By culturing WT PAO1 at a range of temperatures, we found partial inactivation at temperatures higher than 40.5 °C and full inactivation at temperatures above 42 °C (Fig. S1A). Additionally, we determined the duration required for complete iREN to occur at 43 °C to be a minimum of 3 hours of cell division during the early log phase (Fig. 1C). iREN does not occur with transient heat exposure or long-term exposure during the stationary phase (Fig. S1B). This suggests that cell division and *de novo* DNA synthesis are essential for iREN.

Recombination processes, both RecA-dependent and independent, are typically activated in response to DNA damage^16^. To determine if the iREN phenotype depends on these processes, we created mutants in the *rec* genes. Deletion of *recA*, *recB*, and *recC* had no impact on the iREN phenotype (Fig. S2A). Additionally, exposure to UV light or agents causing double-strand DNA breaks, such as fluoroquinolones, did not trigger complete iREN (Fig. S2B). These findings suggest that the iREN phenotype is not dependent on recombination or the SOS response, which was previously implicated in *E. coli* restriction alleviation (RA)^9^. This, along with the clear need for cell division during high-temperature exposure (compared to a transient heat shock), demonstrates that iREN is distinct from RA.

### iREN phenotype heritability

To assess the duration of iREN, we passaged PAO1 cultured at 43 °C for multiple generations at 37 °C while quantifying EOP and plating for colony-forming units (CFUs) to track cell doublings. Restriction activity in the population did not fully recover until around 60 generations of cell division, with partial recovery detectable after the first ∼25 generations (Fig. 2A). Interestingly, after approximately 60 generations of restriction recovery, iREN can occur again by culturing the bacteria at 43 °C (Fig. S2C). This indicates that iREN is not due to a permanent genetic change.

**Figure 2:**
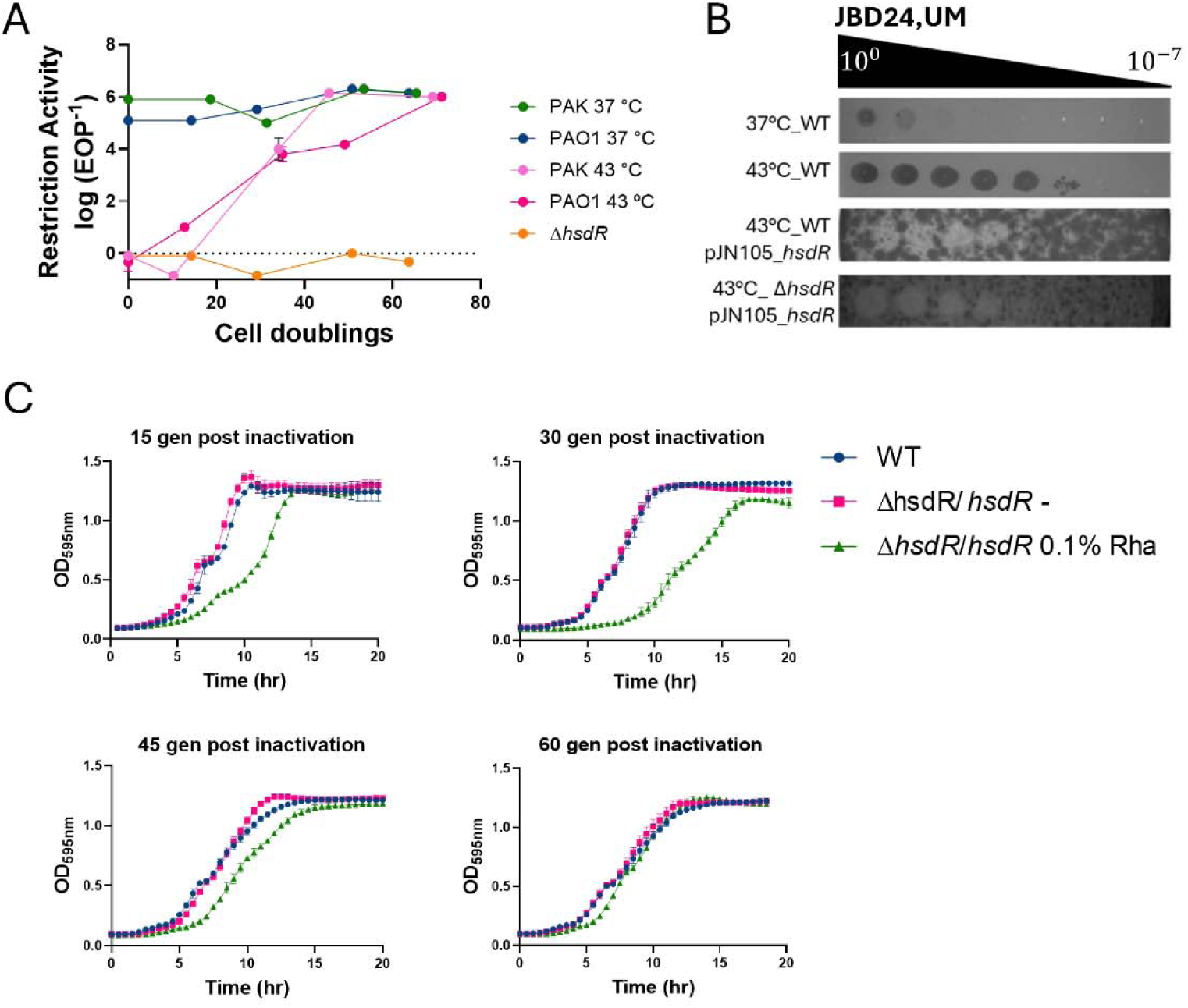
IREN phenotype is heritable and lasts for multiple generations. **(A)** Recovery of restriction activity over generations at 37°C following overnight growth of WT PAO1 and PAK strains at either 43 °C or 37 °C with Δ*hsdR* as a control. **(B)** Plaquing efficiency comparison of strains with or without *hsdR* OE grown at either 37 °C or 43 °C, including WT and *hsdM* mutant strains, when infected with UM phages. **(C)** Restriction forced activation following 43 °C: Growth curve at 37 °C after ON growth at 43 °C of WT (blue) and strains harboring an inducible copy of *hsdR*, with (green) or without (pink) inducer. The inducer was added at t=0. Generations were counted for the WT strain. The graphs in (A) and (C) represent the average of three independent biological replicates, with error bars indicating standard deviation.

To determine whether an inability to inactivate restriction would lead to cell toxicity, we over-expressed HsdR. Interestingly, HsdR over-expression was highly toxic, but only in cells previously grown at 43 °C (Fig. 2B). Conversely, cells grown at 37 °C could tolerate higher levels of HsdR as could PAO1 Δ*hsdM* previously grown at 43 °C (Fig. S4). This demonstrates that inactivation of HsdR is required for cell fitness after growth at 43 °C and that the observed HsdR-induced toxicity is likely due to nucleolytic activity by HsdMSR on unmethylated sites in the bacterial chromosome.

To examine whether restriction activity can be restored sooner than 60 generations, we created a tightly controlled HsdR expression strain in the background of Δ*hsdR* with no promoter leakiness, verified by assessing restriction activity at 37 °C (Fig. S4). We induced HsdR expression at different time points during recovery of restriction. The results showed that inducible re-activation of restriction remains toxic for over ∼45 generations of growth at 37 °C (Fig. 2C). These results demonstrate that bacteria exhibit a memory-like ability to keep HsdR off for multiple generations after growth at 43 °C, which is essential for bacterial fitness. This inactivation of HsdR, however, can be bypassed by HsdR over-expression.

The motifs that Type I R-M homologs target vary based on sequence divergence in the HsdS proteins^17^. To determine whether the iREN phenotype is conserved in different strains of *P. aeruginosa*, we tested the *P. aeruginosa* strain PAK, which restricts JBD24 phages produced by either PAO1 or PA14. We found that PAK also exhibits heritable iREN (Fig. 2A). We further infected distinct clinical and environmental strains of *P. aeruginosa* with UM phage JBD30 and demonstrated that although the restriction systems are highly diverse, these strains also become more phage sensitive following growth at 43 °C^18^ (Table S1).

### Impaired modification activity during 43 °C growth

Since toxicity caused by the forced restriction re-activation indicated HsdMS-dependent self-targeting, we hypothesize that unmodified sites are generated in the bacterial chromosome during 43 °C growth. We first examined the stability of the methyltransferase proteins at elevated temperature conditions by immunoblotting for HsdM and HsdS via epitope tags fused to each protein in separate strains at the native genomic locus. The results clearly show that both HsdM and HsdS are highly unstable during 43 °C growth, as the protein levels were reduced compared to the levels observed at 37 °C (Fig. 3A).

**Figure 3:**
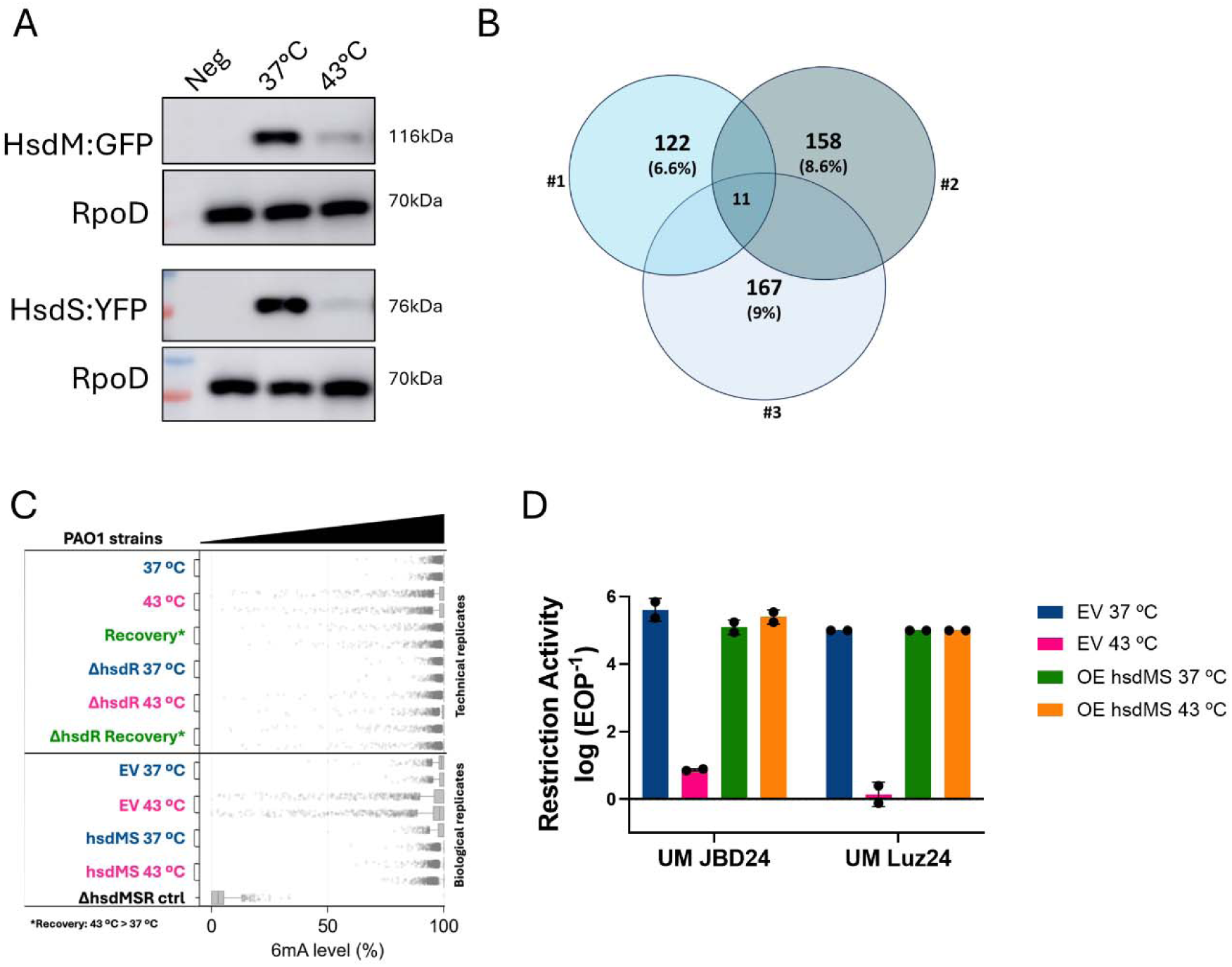
The IREN phenotype arises due to hypomodification during elevated temperature growth conditions. **(A)** Stability of modification proteins under different conditions: Western blot analysis of GFP/YFP-fusion native proteins following growth at different conditions. The presented blot is representative of one of three independent biological replicates, with RpoD as a loading ctrl. **(B-C)** Methylome analysis post-HS: (B) SMRT-seq results for WT strain grown at 43 °C compared to 37 °C. The numbers indicate the count of unmodified sites. Each circle represents one of three independent biological replicates, with shared groups representing conserved sites. (C) e-TAM analysis under different conditions: strains grown at 37 °C, 43 °C, and after initial recovery from 43 °C, single dots represent a possible modification site. Technical and biological replicates illustrate the methylation status. **(D)** OE of HsdMS at 43 °C inhibits iREN: Restriction activity of strains with empty vector and HsdMS OE initially grown at 37 °C (blue and green) or 43 °C (pink and orange), with induction at initial growth followed by sub-culturing without induction at 37 °C. Activity was measured following 10 generations at 37 °C. For (E) graphs, each dot represents an independent biological replicate.

Next, to validate that the decreased protein stability results in methylation deficiency, we directly examined the influence of 43 °C growth on the bacterial chromosome methylation state. For that, we used single-molecule-real-time sequencing (SMRT seq) to analyze the bacterial methylome. Surprisingly, the SMRT seq analysis revealed that while hypomethylation indeed emerges after 43 °C growth, only approximately 8% of the modification sites appeared to be unmodified in the population (Fig. 3B), with 11 shared sites between biological replicates, indicating possible hotspots for deficient methylation.

Given the subtle methylation loss and the complexity of the population, we employed an orthogonal method with higher sensitivity and quantitative abilities, eTAM-seq, to more accurately measure genomic methylation. This method deploys enzymatic modification of genomic DNA to distinguish modified adenines from unmodified adenines via Illumina sequencing^19^. This method similarly revealed a 3% decline in the general methylation of the bacterial genome after 43 °C growth (Fig. 3C). Moreover, decreased methylation percentage is also observed in the recovered population after 10 cell doublings at 37 °C. Remarkably, despite the eTAM-seq and SMRT-seq experiments being executed in independent laboratories and sequenced with distinct methodologies, 10 out of the 11 hypomethylated sites were shared between the two approaches. While it is currently unclear what is unique about these sites that makes them prone to becoming hypomethylated, these data together demonstrate that a surprisingly subtle decrease in genomic methylation explains why HsdR must be turned off and why its forced expression is toxic in an HsdM-dependent manner.

To examine whether loss of methylation activity at 43 °C is necessary to induce loss of HsdR activity, HsdMS were over-expressed during growth at 43 °C. Expression of these proteins during the overnight growth at 43 °C was indeed sufficient to restore genomic methylation and abolish iREN (Fig. 3D), confirming that iREN is driven by methylation deficiency. This was further verified by eTAM-seq results, which showed that genomic methylation remained intact when HsdMS was over-expressed (Fig. 3C) Taken together we propose that cell division at 43 °C destabilizes HsdMS, reduces functional methyltransferase levels, leading to incomplete methylation of de novo produced host DNA, which in turn drives the need for iREN. This genomic hypomethylation then recovers slowly over multiple generations, while HsdR must remain inactive.

### Post-translational decrease in HsdR protein levels during iREN

In PAO1, *hsdM* and *hsdS* are adjacent genes and presumably co-transcribed, while *hsdR* is located within an adjacent operon, suggesting the possibility of independent transcriptional or translational regulation. To gain a better understanding of the different *hsd* components regulation and to find related factors, we performed both transcriptomics and proteomics analyses following five generations after 43 °C growth and at the end of the restriction recovery period (Fig. S6A and 4A). This specific time point was chosen because it aligns with the observed iREN phenotype, allowing us to focus on the effects of the restriction while minimizing effects from high temperature growth. The results showed that at the transcript level, all three genes showed no significant change comparing cells grown only at 37 °C to those five generations after 43 °C growth (Fig. S6A). At the protein level, we also observed that HsdM and HsdS were not decreased five generations after 43 °C growth. However, according to proteomics measurements, HsdR protein levels were significantly reduced after 43 °C growth and eventually returned to basal levels by the end of the recovery phase (Fig. 4A). To validate that HsdR is post-translationally regulated, we performed reverse transcriptase quantitative PCR analysis that showed no significant change in the transcript levels of the *hsd* genes (Fig. S6B). We further analyzed both a transcriptional reporter, a strain carrying *mCherry* instead of the *hsdR* ORF, and a translational reporter, a strain with fused *hsdR*-*mCherry* in the native genomic context (fused to the full-length ORF of *hsdR*). The reporters were analyzed following either 37 °C or 43 °C growth and showed that only the translational reporter exhibited reduced fluorescence after 43 °C growth (Fig. S6C), further corroborating that the HsdR protein is controlled post-translationally.

**Figure 4:**
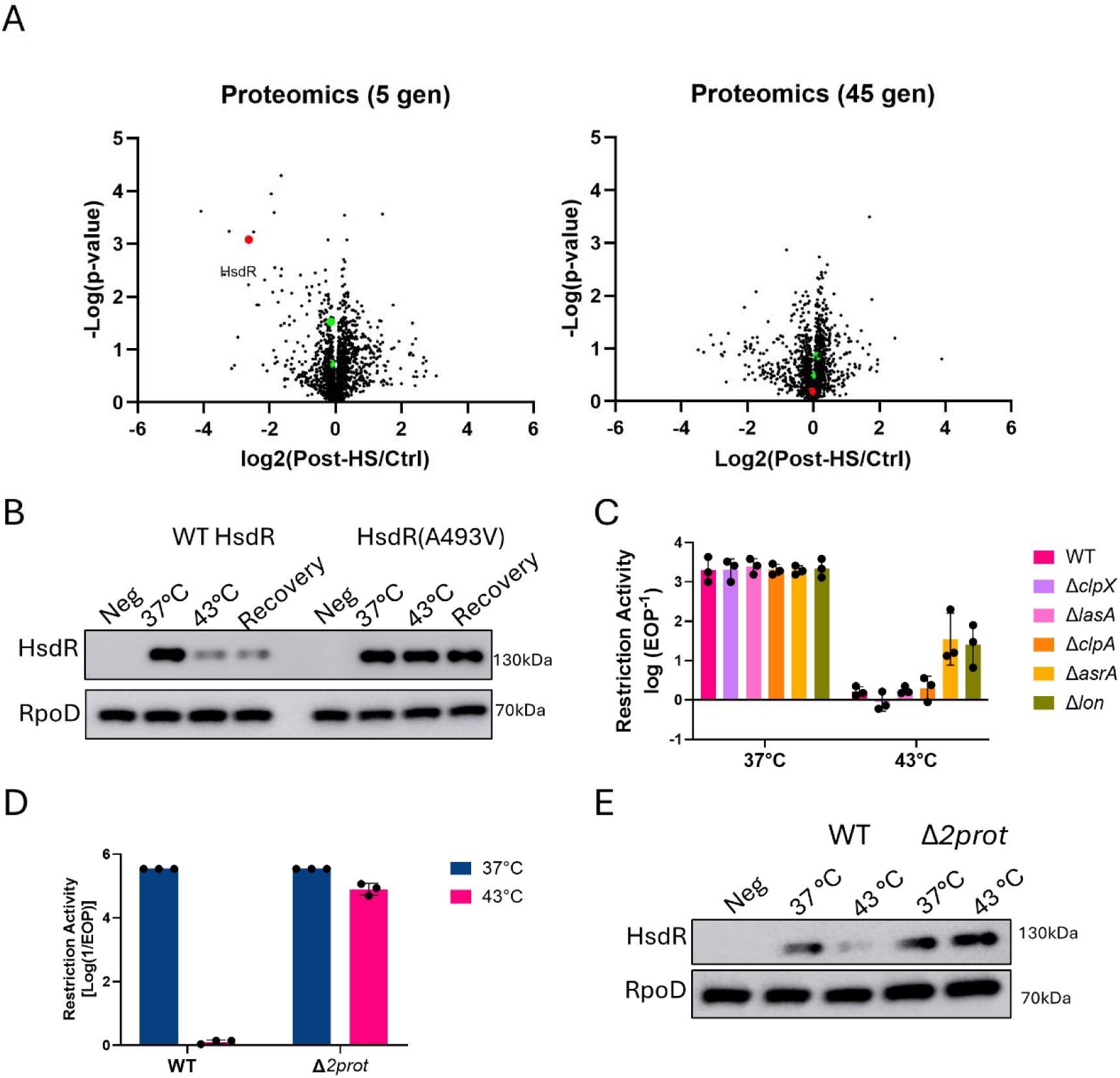
The restriction endonuclease levels decrease in a Lon-like protease-dependent manner. **(A)** Global proteomics analysis: Samples taken five generations following 43 °C growth and at the end of the recovery phase (45 generations) were compared to normal growth conditions. Marked dots indicate Hsd proteins: HsdR (red) and HsdMS (green). **(B)** Decay of HsdR protein levels with translocation activity dependence: Western blots of natively expressed HsdR:flag in WT or translocation-deficient mutant (HsdR(A493V)) strains grown at 37°C, 43°C, or sub-cultured to 37°C following ON growth at 43 °C for 5 generations. **(C)** Proteases effect on iREN: Panel of protease mutants: *clpX* (purple), *lasA* (light-pink), *clpA* (orange), *asrA* (yellow), and *lon* (green). Mutant restriction activity was measured at the end of ON growth at 37°C (left) or 43°C (right). **(D-E**) Regulation of iREN by Lon-like proteases: (D) Restriction activity following ON growth at 37 °C or 43 °C of the double proteases mutant strain compared to WT. (E) HsdR:flag native protein levels at 37°C and post-HS Δ*2prot* compared to WT. For graphs (C) and (D), each dot represents a single biological replicate, with error bars indicating standard deviation. RpoD was used as loading ctrl.

To track HsdR protein levels, we introduced a FLAG-tag (DYKDDDDK) fusion to *hsdR* in its native genomic locus. Western blot analysis shows a reduction in HsdR protein levels both after culturing at 43 °C and after five generations of recovery at 37 °C (Fig. 4B). We additionally immunoblotted a WT PAO1 strain with an HsdR-specific custom antibody which showed a similar reduction in HsdR levels (Fig. S7). For restriction alleviation previously characterized in *E. coli*, it was shown that the DNA translocation activity of the endonuclease is required for proteolytic control^20^. To assess whether this is also the case for iREN, we introduced a mutation in *hsdR* that is defective in ATP hydrolysis and thus cannot perform DNA translocation (HsdR(A493V)). Western blot analysis showed that the ATPase mutant protein levels stayed intact after 43 °C growth (Fig. 4B). These results indicate that either the translocation is essential for protein regulation or that restriction-competent HsdR triggers the loss of HsdR protein.

### Lon-like proteases required for iREN

As shown above, the iREN phenotype is conserved among *P. aeruginosa* strains, so we sought to examine the involvement in iREN of known conserved proteases of *P. aeruginosa*. We generated PAO1 complete ORF deletion mutants of the following proteases: ClpX, LasA, ClpA, AsrA, and Lon. We assessed the restriction activity for the protease mutants at 37 °C and following 43 °C growth. In all protease knockouts, iREN was still observed, however both AsrA and Lon knockouts showed only partial inactivation of the restriction activity (Fig. 4C).

AsrA is a Lon-like protease, highly similar (40% a.a. sequence identity) to Lon^21^. Both are highly conserved and important proteases of *P. aeruginosa*^22,21^. Considering the high similarity between the proteases, we assumed that the partial effect is due to functional compensation. Therefore, we created a double mutant strain lacking both the *asrA* and *lon* genes (Δ*2prot*). Indeed, the restriction activity of the Δ*2prot* strain after growth at 43 °C was almost completely intact (Fig 4D), demonstrating the necessity of these two proteases in iREN. Single and double plasmid complementation restored iREN in the Δ*2prot* background (Fig. S8). With the proteases absent, HsdR protein levels were high after incubation at both 37 °C and 43 °C (Fig. 4E). Taken together, the results indicate that Lon and AsrA are necessary components regulating the proteolysis of HsdR.

### Fitness cost for iREN deficient strain

Forced expression of HsdR was lethal in 43 °C-grown strain due to the presence of unmodified methylation sites in the bacterial chromosome, which led to self-targeting. This raised the question of how the Δ*2prot* strain, lacking both Lon-like proteases, was viable and survived without inactivating restriction. To query whether a more subtle defect emerges when cells cannot proteolyze HsdR, we co-cultured Δ*2prot* cells with Δ*2prot*Δ*hsdR* cells. The competition results showed that the Δ*2prot*Δ*hsdR* outcompeted the Δ*2prot* strain after six hours of growth at 43 °C, whereas the Δ*hsdR* strain showed no advantage over WT under 43 °C growth (Fig. 5A and S9). These results suggest that there is indeed a fitness cost to maintaining an intact Type I R-M system when the regulatory proteases are absent. However, despite this detected defect, the Δ*2prot* strain could still grow at 43° C. One possible explanation for the relatively minimal fitness cost of retaining the Type I R-M system without regulatory proteases is that the HsdMS methyltransferase complex may also be stabilized in the absence of proteolysis.

**Figure 5:**
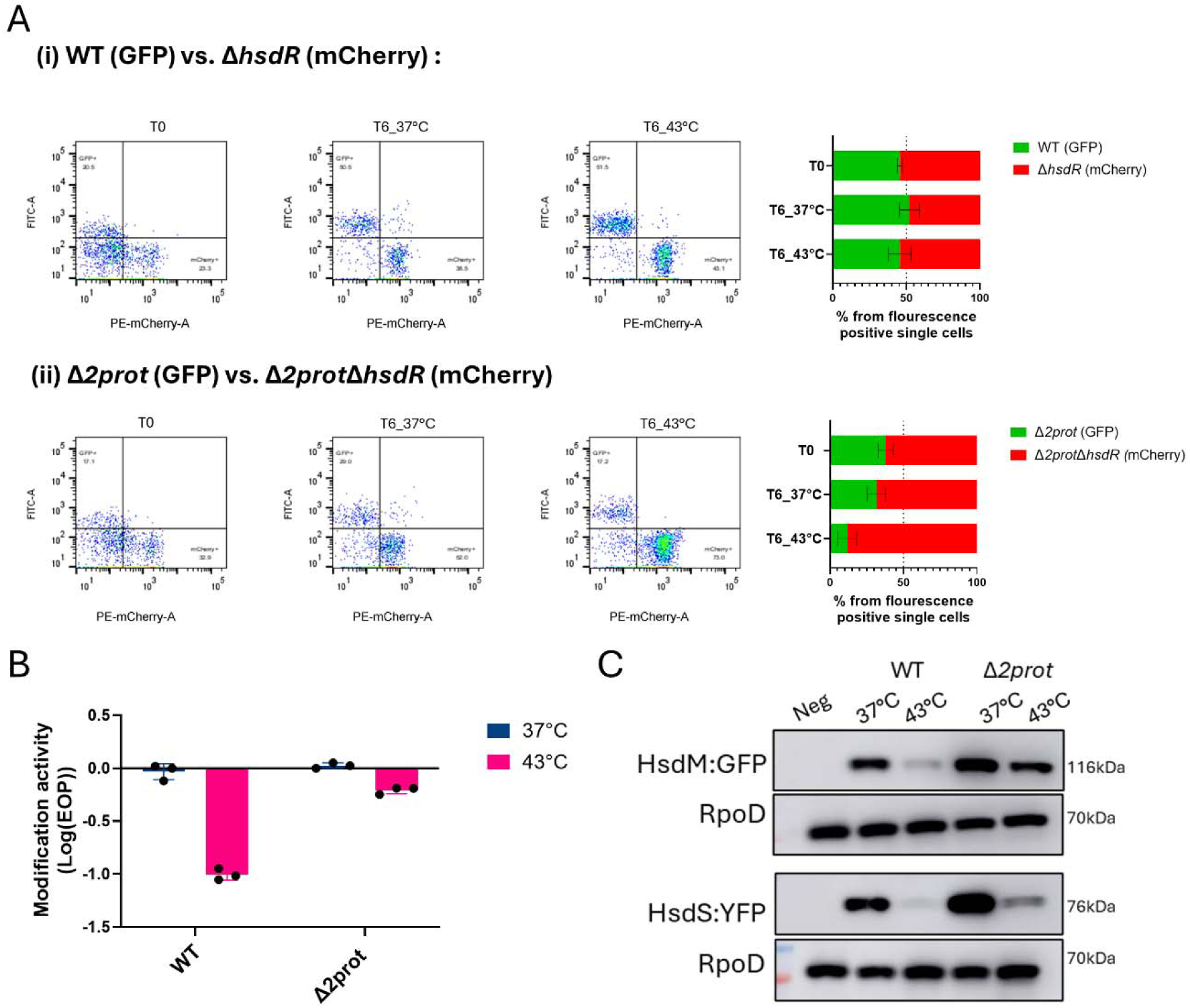
Proteases mutants that cannot degrade HsdR are outcompeted. **(A)** *hsdR* deletion competition under different conditions: Flow-cytometry analysis of population variation in mixed WT and Δ*hsdR* strains (top), or double-proteases mutant and double mutant lacking *hsdR* (bottom). Mixtures were grown for six hours before analysis. Analysis histograms are displayed for representative replicates in both conditions (left), and population percentages are shown as an average of three biological replicates (right). **(B)** Modification activity in protease mutants: Phage modification activity assay for WT and double-protease mutant Δ*2prot* at 37 °C and 43 °C. Each dot represents a single biological replicate. **(C)** Modification proteins stability in protease mutants: Western blot analysis of GFP/YFP-fusion native proteins following growth at different conditions in Δ*2prot* strain compared to WT. The presented blot is representative of one of three independent biological replicates. RpoD was used as loading ctrl.

For a simple assay to query how active modification is in the cell, we assessed the degree of modification of produced phages. Specifically, we examined whether phages produced from cells grown at 43 °C were modified by comparing their titer on WT PAO1 to Δ*hsdR*. Indeed, for the WT background, approximately 90% of the phage population produced by cells at 43 °C was unmodified, as evidenced by a ten-fold reduction in EOP when plated on restrictive WT PAO1. This assay effectively detected the modification deficiency in WT cells grown at elevated temperatures. The modification activity assay, based on phage genome modification, confirmed that most phages produced by the Δ*2prot* strain were modified (Fig. 5B). Moreover, HsdMS methyltransferase levels were more stable in the Δ*2prot* strain at 43 °C (Fig. 5C). Together these findings suggest that Lon proteases reduce the stability of both modification proteins and HsdR during high-temperature growth, which explains why the Δ*2prot* strain does not exhibit a severe growth defect: because HsdMS and methyltransferase activities are largely intact.

### Proteases specificity in iREN

This work has implicated two Lon proteases in the decrease of three Type I R-M proteins (HsdMSR) during cell division at temperatures >41 °C. This raises the final question as to which of the R-M proteins are targeted by each protease. We therefore assessed protein levels during initial growth at 43 °C compared to 37 °C in different protease mutant backgrounds.

The results indicate that the effect of the proteases are differentiated. Lon protease mutant resulted mainly in elevated HsdR protein levels following growth at 43 °C, while deletion of AsrA protease resulted in increased HsdM stability during initial growth at 43 °C. Finally, for HsdS, an additive effect was observed as the single protease mutants showed only a slight increase in protein levels, suggesting that both proteases act on HsdS (Fig. 6).

**Figure 6:**
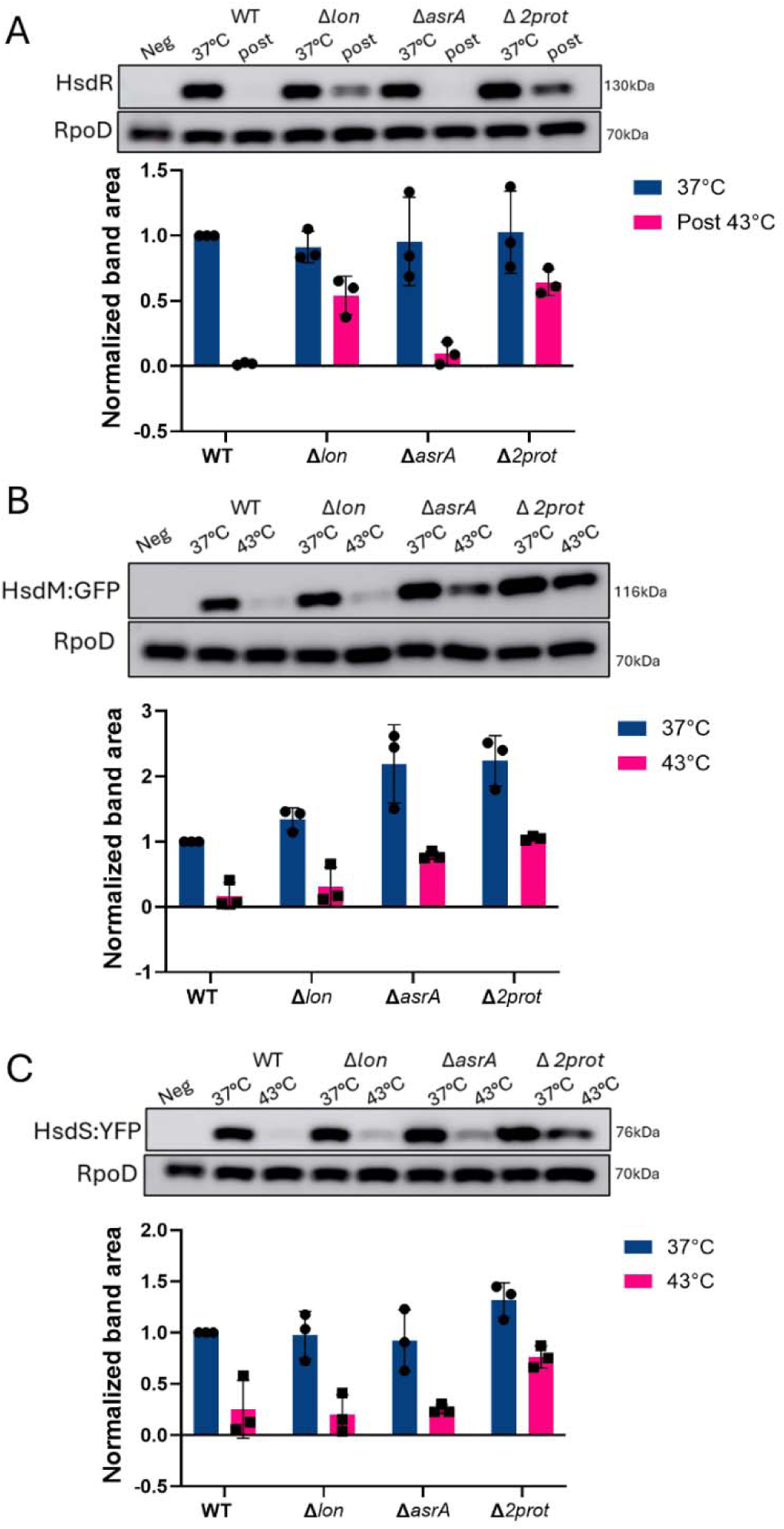
Lon-like proteases operate in iREN in collaborative manner. **(A)** HsdR protein levels in protease mutants: Western blot analysis of flag-fusion native HsdR protein following growth under different conditions of WT and protease mutants. The quantification of band area is normalized to the loading control (RpoD) and WT at 37 °C as a standard (bottom panel). **(B-C)** Stability of modification proteins in protease mutants: Western blot analysis of GFP/YFP-fusion native HsdM (B) and HsdS (C) proteins following initial growth under different conditions, with quantification of band area normalized to the loading control (RpoD) and WT at 37 °C as a standard (bottom panels). The blot presented is representative of one of three independent biological replicates.

These findings suggest a complex regulatory mechanism in which Lon and AsrA proteases collaboratively down-regulate the levels of HsdR, HsdM, and HsdS proteins at 37 °C and 43 °C. These activities are likely required for the genetic maintenance of Type I R-M systems by inactivating the endonuclease holoenzyme when genomic hypomethylation emerges.

## Discussion

Holloway first described restriction inactivation at high temperatures, identifying a heritable restriction loss phenotype under such conditions ^15^. Our findings identified the inactivated type I R-M system, demonstrated that this is a common property of *P. aeruginosa* isolates, and implicated subtle hypomethylation in the genome as the driver. To this end, we identified that only a minority of genomic methylation sites become unmodified during high-temperature growth. Moreover, we found that both hypomethylation and the iREN phenotype are proteolytically regulated by Lon-like proteases in a collaborative manner (Fig. 7).

**Figure 7:**
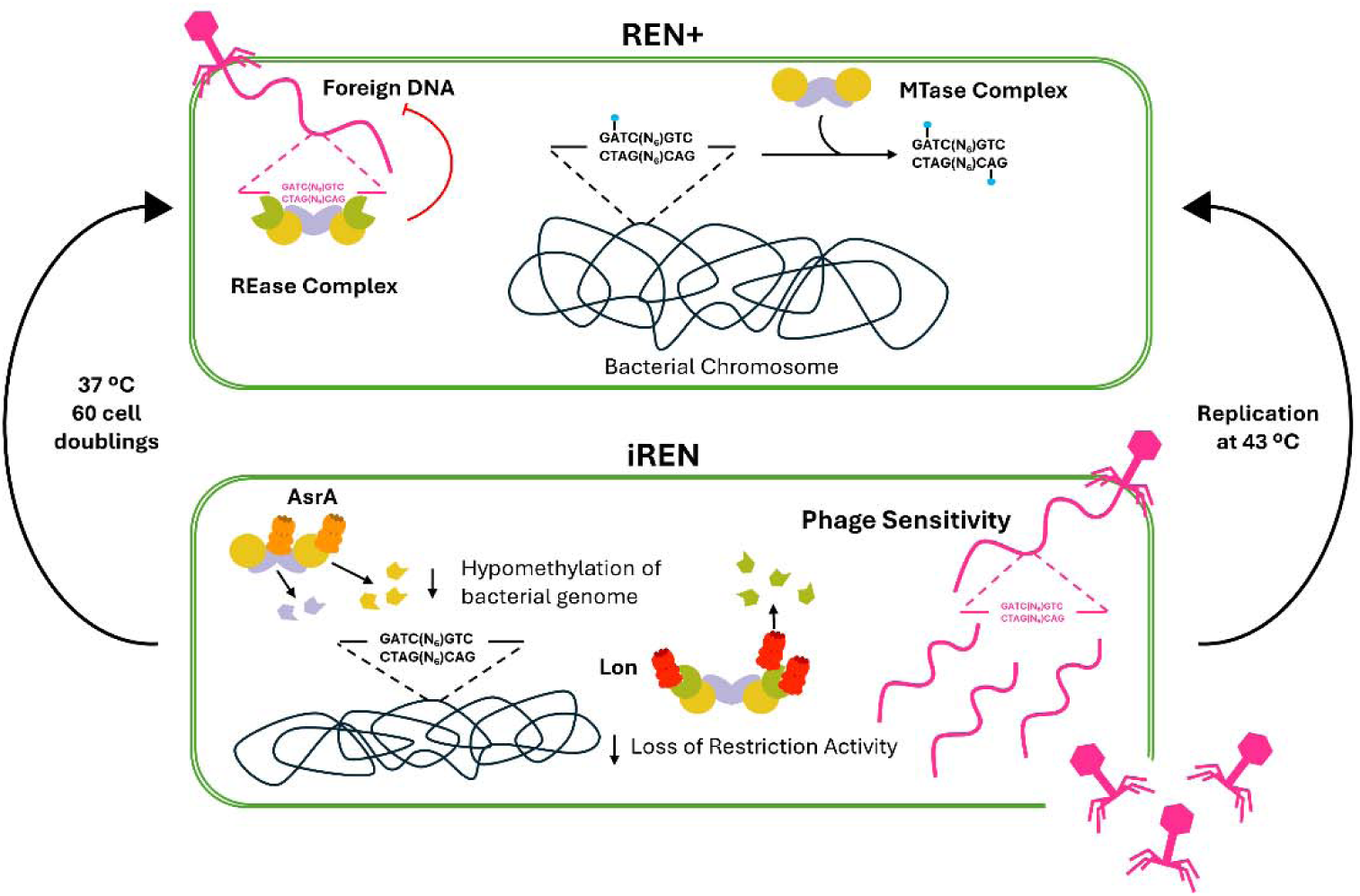
Proposed model for iREN. Under normal conditions (top, **<u>R</u>**estriction **<u>EN</u>**donuclease **+** or **REN+**), PAO1 maintains a functional type I R-M system, with the MTase complex (HsdM in yellow, HsdS in grey) modifying the host genome and the REase complex (including HsdR in green) restricting foreign DNA. After growth at 43 °C (bottom, **iREN**), the restriction targets unmodified sites on the bacterial chromosome, which is inhibited by Lon-like proteases (red triangle). Following approximately 60 generations of recovery at 37 °C, bacteria restore their native restriction activity.

iREN exhibits a novel type of temporal restriction loss of the type I R-M system when the bacterial genome becomes undermethylated. RA in *E. coli* is induced upon acquiring a new type I R-M system and exposure to UV and other DNA-damaging agents. Recently, it was also shown to be activated through plasmid-carrying hemi-modified recognition sites^9^. Unlike RA, we found that iREN for *P. aeruginosa* is not induced upon DNA breaks and is a *recA* and *recBC*-independent process. This suggests that iREN is closely associated with the instability of HsdMS, as stable HsdMS in *P. aeruginosa* sustain normal methylation rates even when recombination processes create hemi-modified sites. The recovery from iREN took up to 60 generations, starting with only 3-8% of unmodified sites in the genome. In contrast, for RA, when acquiring a new system with a fully unmodified chromosome, only a lag of 15 generations was described^7^ indicating either a faster modification rate of the *EcoKI* system or, alternatively, a tighter regulatory mechanism for iREN.

The observed high sensitivity to complete restriction inhibition, even with only a slight decrease in methylation, highlights the critical need for strict autoimmunity regulation in bacterial defense systems, as nearly all bacterial defense systems possess autotoxicity and must be tightly controlled to prevent unnecessary activation. Systems operating through abortive infection (Abi) mechanisms inhibit cellular metabolism or program bacterial cell dormancy/death upon infection to prevent viral replication. For example, Toxin-Antitoxin (TA)^23^ systems neutralize toxins via labile antitoxins until changes in cellular physiology license toxin activity. Direct defense systems, such as R-M and CRISPR-Cas^24^ systems, also exhibit autotoxicity from off-target nucleolytic cleavage of host DNA or deficiencies in protective mechanisms. While our findings focused on type I R-M, this temperature-induced restriction loss serves as a valuable model for understanding how genomic methylation may be compromised and responded to. Moreover, based on our results, these mechanisms presumably vary between bacterial species and should be investigated in other hosts.

Our findings suggest that Lon proteases have a dual regulatory role: they initially cause hypomethylation during high-temperature growth by reducing modification activity, but also degrade HsdR to limit generation of toxic double-stranded breaks in the bacterial chromosome and provide time for restored methylation. The restriction recovery process itself was found to be highly reproducible but gradual, consistently occurring after a specific number of bacterial cell divisions. This moderated recovery could result from population heterogeneity, where a subset of cells fully recovers and eventually dominates the culture over 60 generations, or from chromosomal re-modification, where the proteases degrade the restriction component, leading to partial inhibition and reflecting the chromosomal methylation status. Understanding these mechanisms will require further studies at the single-cell level.

These insights have practical implications in laboratory settings. iREN represents a rapid and simple experimental perturbation to bypass a common bacterial defense, which is particularly important when working with phages that infect multiple related strains. Given the observed memory phenotype of iREN, understanding its effects on the host is essential to manage and control its application in laboratory experiments. In conclusion, this study reveals the importance of precise negative regulation of a bacterial defense system to maintain genomic integrity with memory to ensure safety. Our findings highlight how bacteria quickly adapt to methylation deficiencies, using high-temperature growth as a model for understanding these responses.

## Author contributions statement

E.S., A.V., S.D.M., and I.A. generated strains and conducted phenotypic experiments and analyses. E.S. drafted the initial manuscript, and A.V., S.S., E.B., and J.B.D. reviewed and edited the text. Y.W. and H.Y. prepared the eTAM-seq libraries, while C.Y. performed the next-generation sequencing and bioinformatics analysis, with additional analysis contributions from I.L.L. W.T. supervised the eTAM-seq work. S.L. contributed to the methodology. E.B. and J.B.D. provided overall supervision and secured funding.

## Acknowledgments

This work is part of the Ph.D. thesis of E.S. Partial funding for this work was through the Dyna and Fala Weinstock Foundation to E.B., the President’s Scholarships, and the Merit-Based Scholarships at the Institute of Nanotechnology of Bar-Ilan University for E.S. Experiments were funded by awards to J.B.D. by the Bowes Biomedical Investigator Award and the UCSF Sanghvi-Agarwal Innovation Award research gifts. A.V. was supported by NSF and S.D.M. by an NIH F31 award. We thank The Crown Genomics Institute of the Nancy and Stephen Grand Israel National Center for Personalized Medicine and the Weizmann Institute of Science for proteomic data and SMRT-seq processing and analysis support. We thank Rob Lavigne for sharing the Luz24 phage. The eTAM-seq section is supported by a pilot award under grant no. RM1 HG008935. We thank Dr. Qing Dai for constructive suggestions on eTAM-seq library preparation. We thank the Single Cell Immunophenotyping Core Facility at the University of Chicago for sequencing support.

## Data availability

All strains and plasmids utilized in this study are detailed in Table S2. The raw data from SMRT-seq, RNA-seq, and liquid chromatography-mass spectrometry have been deposited and are available at: https://doi.org/10.25452/figshare.plus.c.7498956.

## Materials and Methods

### Bacterial strains, plasmids, and phages

The bacterial strains, phages, and plasmids used in this study are detailed in Supplementary Table S2. Unless stated otherwise, all strains were grown in LB (Luria-Bertani broth, Difco) at 37 °C. For deletion mutant creation, the following media were used: Vogel Bonner Minimal Medium (VBMM)^25^, Pseudomonas Isolation Agar (PIA, Difco), and No salt Luria-Bertani (NSLB) supplemented with 10% sucrose. For DH5α heat shock, BHI (brain heart infusion broth, Difco) media was used. All strains were grown at 37 °C unless otherwise specified. Antibiotic concentrations used in this study were 300 μg/mL Carbenicillin (Crb) and 50 μg/ml Gentamicin (Gm) for *P. aeruginosa*, 100 μg/mL Ampicillin (Amp), and 30 μg/mL Gm for *Escherichia coli*.

### Plasmids construction

The genomic extraction was carried out using the DNeasy Blood& Cell Culture DNA Kit (Qiagen). For DNA fragment amplification, Phusion® High-Fidelity DNA Polymerase (Thermo) was used. For gene overexpression, primers were designed to complement the beginning and end of each gene, with the addition of either enzyme restriction sites for ligation or an overlap sequence for Gibson assembly. The amplified inserts were purified using NucleoSpin® Gel and PCR Clean-up (MACHEREY-NAGEL). For the ligation assay, inserts and plasmids were digested using the appropriate fast-digest restricted enzymes (Thermo). Ligation was conducted using Biogase - fast ligation kit (Bio-Lab Ltd.). For the Gibson assembly, inserts were incubated in the appropriate concentration with a linearized plasmid and 2X LigON mixture (EURX). For plasmid extraction, the QIAprep spin mini-prep kit (QIAGEN) was used. For verification of successful plasmid transformations, the DNA Polymerase ReddyMix™ PCR Kit and universal primers were used.

### Efficiency of plaquing (EOP) assay

For phage extraction, overnight cultures of JBD24 lysogenic bacteria were prepared, with PAO1 for modified and PA14 for UM phages. These cultures were diluted 1:50 with fresh medium and incubated until reaching an optical density (OD) of 0.5 at 595 nm. Phage induction was initiated by adding 0.4 μg/ml Norfloxacin (NOR, Sigma) antibiotic, followed by an additional incubation period of 1 hour. Fresh LB medium was then added, constituting 50% of the culture volume, and the cultures were further incubated for an hour to facilitate phage amplification. Subsequently, the bacterial cells were centrifuged at maximum speed, and the supernatant was filtered using a 0.45 µm filter (Whatman).

For Efficiency of Plaquing (EOP) calculation, overnight cultures of host bacteria, grown at either 37 °C or 43 °C, were used. Soft-agar LB containing 0.5% agar, preheated to 50 °C, was gently mixed with a volume of 100 µl of the overnight host cultures and poured onto the surface of a solidified agar plate containing 1.5% agar. After appropriate air-drying, serial dilutions of the examined phages were plated in drops of 2 µl on top of the host-containing soft agar. The plates were then incubated overnight at 37 °C, allowing plaques to form, which could be subsequently counted for the calculation of plaque-forming units per milliliter (PFU/ml).

### Restriction and Modification activity calculation

Restriction activity for the examined strain was calculated by its EOP and the EOP of permissive Δ*hsdR* strain with the following formula:

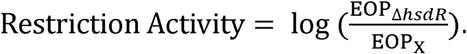

The calculation of modification activity for the examined phage was determined by comparing its EOP in a restrictive strain to the EOP of a permissive Δ*hsdR* strain, using the following formula:

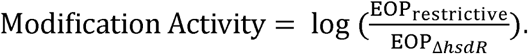

### Growth curve

Overnight cultures of the strains were diluted to 0.005 OD (595 nm) in fresh media and transferred to a 96-well plate, 200μl in each well. Arabinose or Rhamnose was added for gene induction (33.3Mm and 0.2% unless otherwise specified). The plates were incubated for 20 hours at 37 °C or 43 °C with agitation. Optical density measurements at 600 nm were taken every 10 minutes using the Synergy™ 2 Multi-Detection Microplate Reader (BioTek).

### Restriction recovery assay

For the analysis of restriction activity recovery, overnight cultures grown at either 43 °C or 37 °C were assessed for EOP and plating efficiency. These cultures were then sub-cultured by diluting them 1:1000 and incubating them further at 37 °C. EOP and plating efficiency measurements were taken every 12 hours thereafter with repeated sub-culturing passaging of the bacteria. From EOP measurements, restriction activity could be calculated, and from plating efficiency measurements, generations were determined using the following formula:

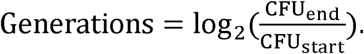

### Protein extraction

Overnight cultures grown at 37 °C or 43 °C were diluted 1:100 into fresh LB and grown at the examined temperature to 0.8 OD (595 nm). From each strain, 8 OD (595 nm) of bacteria were harvested, centrifuged at 4,000 RPM for 10 minutes, and the supernatant was discarded. The cell pellet was then resuspended in lysis buffer (comprising 100 mM NaCl, 5% Glycerol, and 50 mM Tris pH 7.5) supplemented with Benzonase® Endonuclease (MILLIPORE), cOmplete™ protease inhibitor cocktail (ROCHE), and Lysozyme (Sigma-Aldrich). Subsequently, the samples underwent sonication (3 minutes, with cycles of 5 seconds on and 5 seconds off, at 54% amplitude). The sonicated samples were then centrifuged at 14,000 g for 10 minutes, and the upper liquid phase containing the proteins was carefully collected.

### Western blot

For Western blot analysis, protein samples were diluted with Sample buffer (comprising 150 mM Tris-HCl pH=6.8, 3% β-Mercaptoethanol, 6% Sodium dodecyl Sulfate, 0.3% Bromophenol blue, 30% Glycerol, and water), followed by incubation at 95 °C for 10 minutes. Subsequently, the samples were centrifuged at 14,000g for 2 minutes. The processed samples were then separated on a 20% Tris-Glycine gel and transferred onto a nitrocellulose membrane. For blocking, the membrane was treated with 5% skim milk in TBS, and incubated overnight at 4 °C. Following blocking, the membrane was incubated for 1 hour with anti-FLAG antibodies (diluted at 1:2,500, Sigma-Aldrich). After three washes with TBST (Tris-buffered saline with Tween), the membrane was then incubated with mouse anti-mouse (HRP) antibodies (diluted at 1:10,000, Merck) for an hour. Subsequently, the membrane underwent an additional three washes with TBST before being developed using an ECL kit (Thermo).

### Proteomics analysis

For proteomics sample preparation, the samples were subjected to in-solution tryptic digestion using suspension trapping (S-trap method by Protifi). The resulting peptides were analyzed in Liquid chromatography mass spectrometry (LC-MS), using nanoflow liquid chromatography (nanoAcquity) coupled to high resolution, high mass accuracy mass spectrometry (Q-Exactive HF). Each sample was analyzed on the instrument separately in a random order in discovery mode. Raw data was processed with MaxQuant v1.6.6.0. The data was searched with the Andromeda search engine against the *Pseudomonas aeruginosa* PAO1 database as downloaded from Uniprot, appended with common lab protein contaminants. Search parameters included the following modifications: Fixed modification-cysteine carbamidomethylation. Variable modifications-methionine oxidation. The quantitative comparisons were calculated using Perseus v1.6.0.7. Decoy hits were filtered out, and only proteins that were identified in at least two replicates of at least one experimental group were kept.

### Samples collection for proteomics and transcriptomics analysis

Overnight bacterial cultures cultivated at either 37 °C or 43 °C were diluted 1:100 into 15 ml fresh LB medium and cultured until reaching an optical density (OD) of 0.6 at 595nm from each sample, 2 ml was centrifuged and stored at −80°C for future RNA and protein extraction, while the remaining culture was further diluted 1:1000 for overnight growth at 37 °C. This process was repeated daily until the fifth incubation. Finally, RNA extraction was performed simultaneously on samples collected at all specified time points as previously decribed^26^.

### Transcriptomics analysis

For RNA sequencing, 2 ug of total RNA was used for the RiboMinus™ Bacteria Transcriptome Isolation Kit (Invitrogen). The library was constructed with Kapa Stranded RNA-Seq Kit (KK8421) according to the manufacturer’s instructions using 30ng of depleted RNA as starting material. The final quality was evaluated by TapeStation High Sensitivity D1000 Assay (Agilent Technologies, CA, USA). Sequencing was performed based on Qubit values and loaded onto an Illumina MiSeq using the MiSeq V2 (50-cycles) Kit Illumina (CA, USA). Paired-end RNA-seq protocol yielded about 3.4-6.5 million paired-end reads per sample. FastQC (v0.11.2) (https://www.bioinformatics.babraham.ac.uk/projects/fastqc) was used to assess the quality of raw reads. Reads were aligned to *P. aeruginosa* PAO1 strain (assembly GCF_000006765.1) using the bowtie2 aligner software (v2.3.2)^27^ with default parameters. GTF annotation file for the PAO1 strain was downloaded from Pseudomonas Genome DB ( www.pseudomonas.com). Raw read counts for 5708 gene-level features were determined using HTSeq-count^28^ with the intersection-strict mode. Differentially expressed genes were determined with the R Bioconductor package DESeq2^29^ ( Release 3.14). The p-values were corrected with the Benjamini-Hochberg FDR procedure. Genes with adjusted p-values < 0.05 and |log fold change| > 1 were considered as differentially expressed.

### Fluorescence reporter assay

Overnight cultures of bacterial strains containing the *mCherry*-fused reporter from 37 °C or 43°C were sub-cultured to a final concentration of 0.005 OD (595 nm) in fresh media and transferred to a 96-well plate, with 200μL in each well. Following 20 hours of incubation at 37 °C with shaking, optical density at 595 nm, and fluorescence, excited at 580 nm and emitted at 610 nm, were measured using the Synergy™ 2 Multi-Detection Microplate Reader (BioTek).

### Competition assay

Overnight cultures of strains containing plasmids with constitutive expression of either GFP or *mCherry* were diluted to create mixed cultures with a final concentration of 0.025 from each strain. Samples were collected at the initial time point and after six hours of incubation at either 37 °C or 43 °C by centrifugation at max speed, and the supernatant was removed. For fixation, the samples were re-suspended in 1 ml of 4% paraformaldehyde (PFA) for 1.5 hours on ice, followed by a wash with PBS. Samples were also stained with Hoechst (Thermo) for general staining by incubating the samples with 50µl of 1µg/ml concentration. Washed samples was then analyzed by BD LSRFortessa™, results were analyzed by FlowJo™.

### SMRT seq

Pac Bio subreads were mapped to the reference genome *Pseudomonas aeruginosa* PAO1 GCA000006765, ASM676v1, for each sample separately, using the pbmm2 tool from the smrttools 10 toolkit (Pac Bio). 6mA modifications were then inferred with the ipdSummary tools. The control model was used in two separate runs. In the first run, samples from group 43 were used as a reference. In the second run, samples from group 37 °C were used as a reference. Sites with IPD-ratio with p-value < 0.05 were considered to have a new modification in the sample, compared with the reference. Each sample was tested with the reference of the same line, e.g. 1_43_5g was tested against 1_43 or 1_37. 2_43_22g was tested against 2_43 or 2_37, and so on.

### eTAM seq

Genomic DNA was extracted from cells cultured at 37 °C, 43 °C, or after 10 generations of recovery at 37 °C. Cells were 1:1 treated with Lysis Buffer (20mM Tris, 2mM EDTA, 1% SDS, 100µg/mL RNase A, and 100µg/mL Proteinase K) for incubation at 37 °C for 30 minutes, followed by 30 minutes at 55 °C. Following cell lysis, the Zymo Genomic DNA Clean and Concentrator^TM^ kit was used as instructed to extract DNA.

For each DNA sample, 100 ng was fragmented using NEBNext® dsDNA Fragmentase (NEB, catalog no. M0348S) at 37 °C for 20 min and then denatured in 0.1 M NaOH at 37 °C for 5 min. The denatured DNA was neutralized by 10% HOAc, purified by the Oligo Clean & Concentrator kit (Zymo), and mixed with 0.001% (w:w) spike-in probes containing one 6mA site and one inosine site (GTG TCT GGT GTT CTG TCG TGT GCT ACT C/iN6Me-dA/T CCG ATC TCG CAT C/ideoxyI/T CAC AGT ATT CGT CGT ATG AGA CAC AAC TAC ATG CTT GTC CGC TCT TGT GTC GGC T).

The fragmented and denatured DNA was divided into two halves and designated to eTAM-treated and FTO-treated groups. The FTO-treated group was first denatured at 95 °C for 5 min and demethylated by incubating with 200 pmol of FTO and 1× FTO reaction buffer (2 mM sodium ascorbate (Sigma-Aldrich), 65 μM ammonium iron(II) sulfate (Sigma-Aldrich), 0.3 mM α-ketoglutarate (Sigma-Aldrich), 0.1 mg ml^−1^ of bovine serum albumin (NEB) and 50 mM Hepes-KOH, pH 7.0) in reaction volume of 50 ul at 37 °C for 1 h. The demethylated DNA was then purified by the Oligo Clean & Concentrator kit.

Both FTO-treated and eTAM-treated groups were denatured in 10% DMSO at 95 °C for 5 min and deaminated with 200 pmol of TadA8.20 and 1x deamination buffer (50 mM Tris, 25 mM KCl, 2.5 mM MgCl_2_, 2 mM DTT and 10% (v:v) glycerol, pH 7.5) in reaction volume of 10 ul at 44 °C for 3 h and 44 °C - 55 °C with 3-min incubation per degree. The deaminated DNA was denatured by 0.1 M NaOH at 37 °C for 5 min, neutralized by 10% HOAc, and purified by the Oligo Clean & Concentrator kit. The deamination reaction was then repeated at 44 °C for 1 h and 44 °C - 55 °C with 3-min incubation per degree, followed by a final purification using the Oligo Clean & Concentrator kit.

The treated DNA in both groups was then prepared for next-generation sequencing using the xGen™ Methyl-Seq DNA Library Prep Kit (IDT, catalog no. 10009824) following the manufacturer’s directions. The resulting library was purified by AMPure XP beads (Beckman Coulter, catalog no. A63882) following the manufacturer’s directions and submitted for sequencing. After sequencing, paired-end sequencing reads were preprocessed using Cutadapt to remove adapter sequences and polyC tails with the parameters:

“-a AGATCGGAAGAGCACACGTCTGAACTCCAGTCAC -a C{20} -A AGATCGGAAGAGCGTCGTGTAGGGAAAGAG -G G{20}”.

Clean reads were then aligned to the *P. aeruginosa* PAO1 strain (assembly GCF_000006765.1) using the Hisat2-3n^30^ aligner software with default parameters, except for --no-spliced-alignment and --base-change A,G. 6mA sites were detected using the hisat-3n-table command, and the 6mA ratio was calculated as the number of unconverted reads over the total coverage at each A site. Putative 6mA sites were selected based on a negative binomial distribution, adapted from previous methods^31^, and filtered by a sequencing coverage threshold of 20. For systematic comparisons across multiple conditions, all 6mA sites within GATC(N)_6_GTC motifs were extracted for analysis.

---

**Figure S1:**
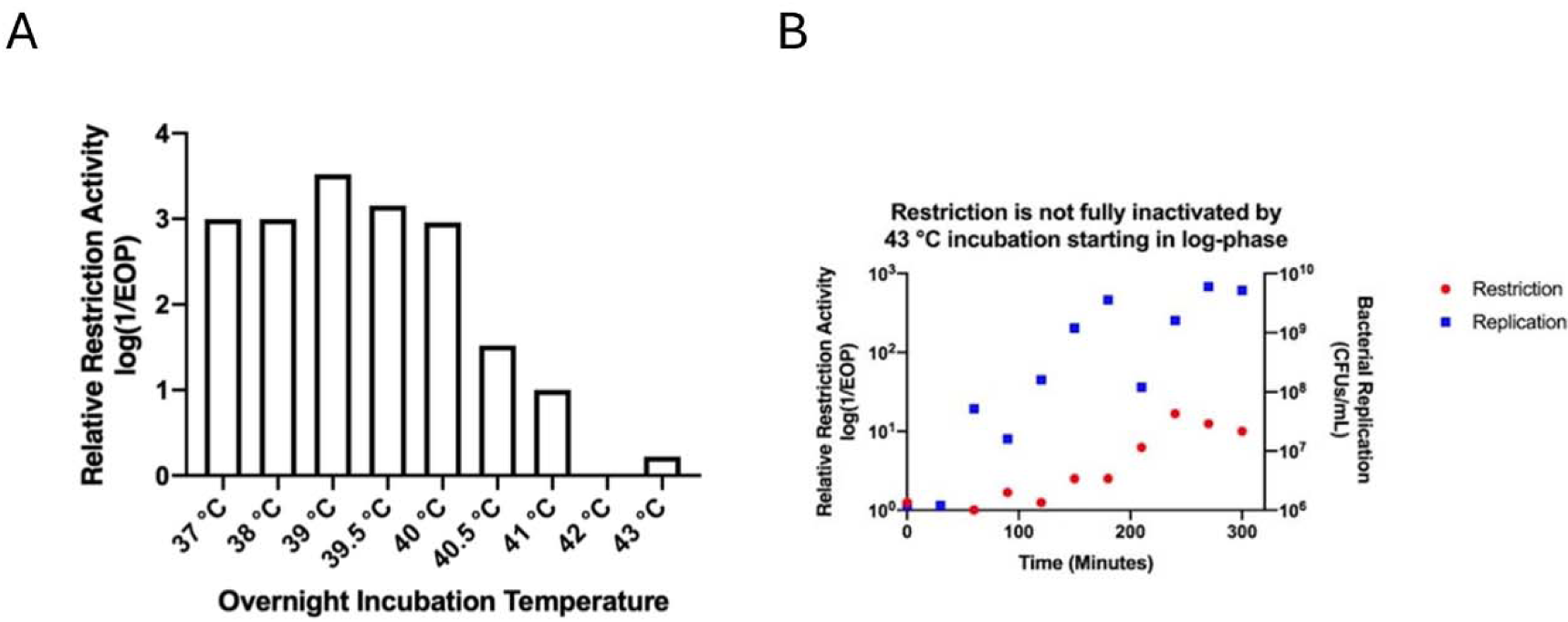
Dynamics and conservancy of iREN. **(A)** Examination of restriction activity following overnight growth at temperatures **(B)** Restriction activity (red) examined for PAO1 WT strain when incubated at 43 °C during different growth stages (blue, OD 600nm).

**Fig. S2:**
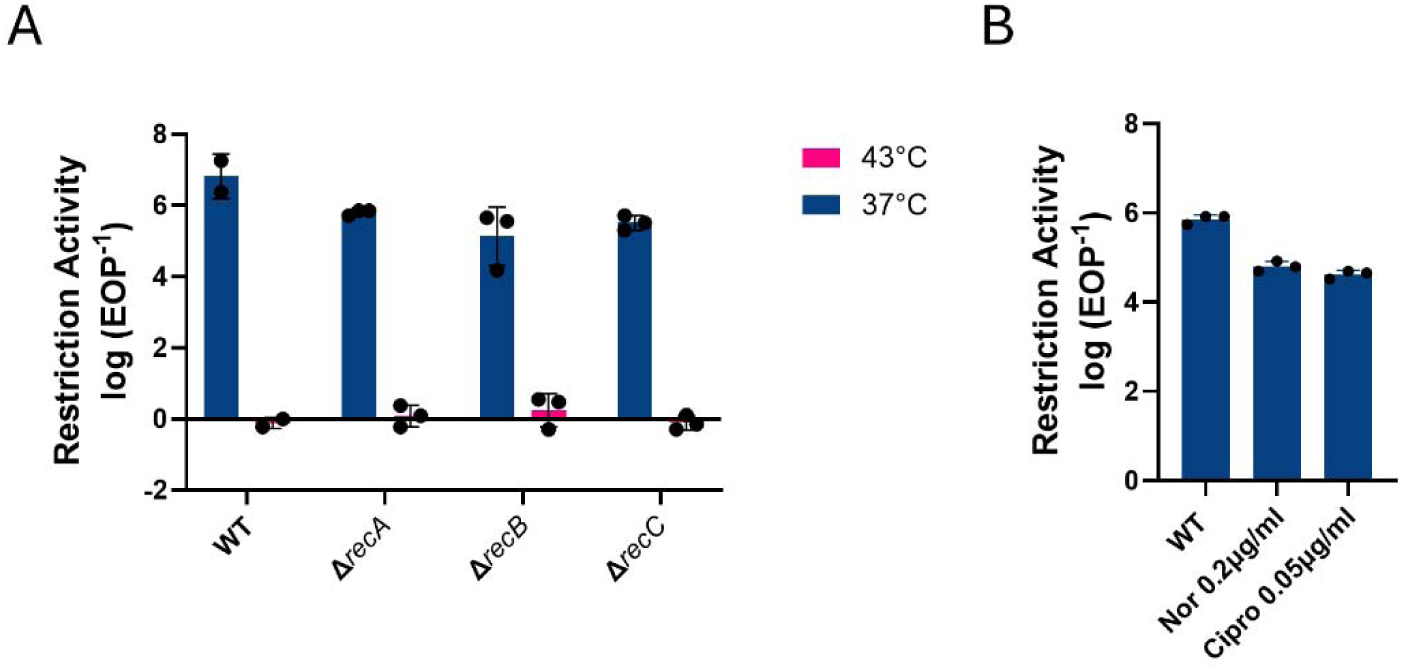
DNA recombination and breaks do not induce iREN. **(A)** restriction activity recombination genes mutant strains at 37 °C (blue) and following 43 °C growth, compared to WT strain. **(B)** Restriction activity following 6 hr growth with sub-inhibitory concentration Fluoroquinolone treatment of Ciprofloxacin and Norfloxacin, compared to untreated WT strain.

**Fig. S3:**
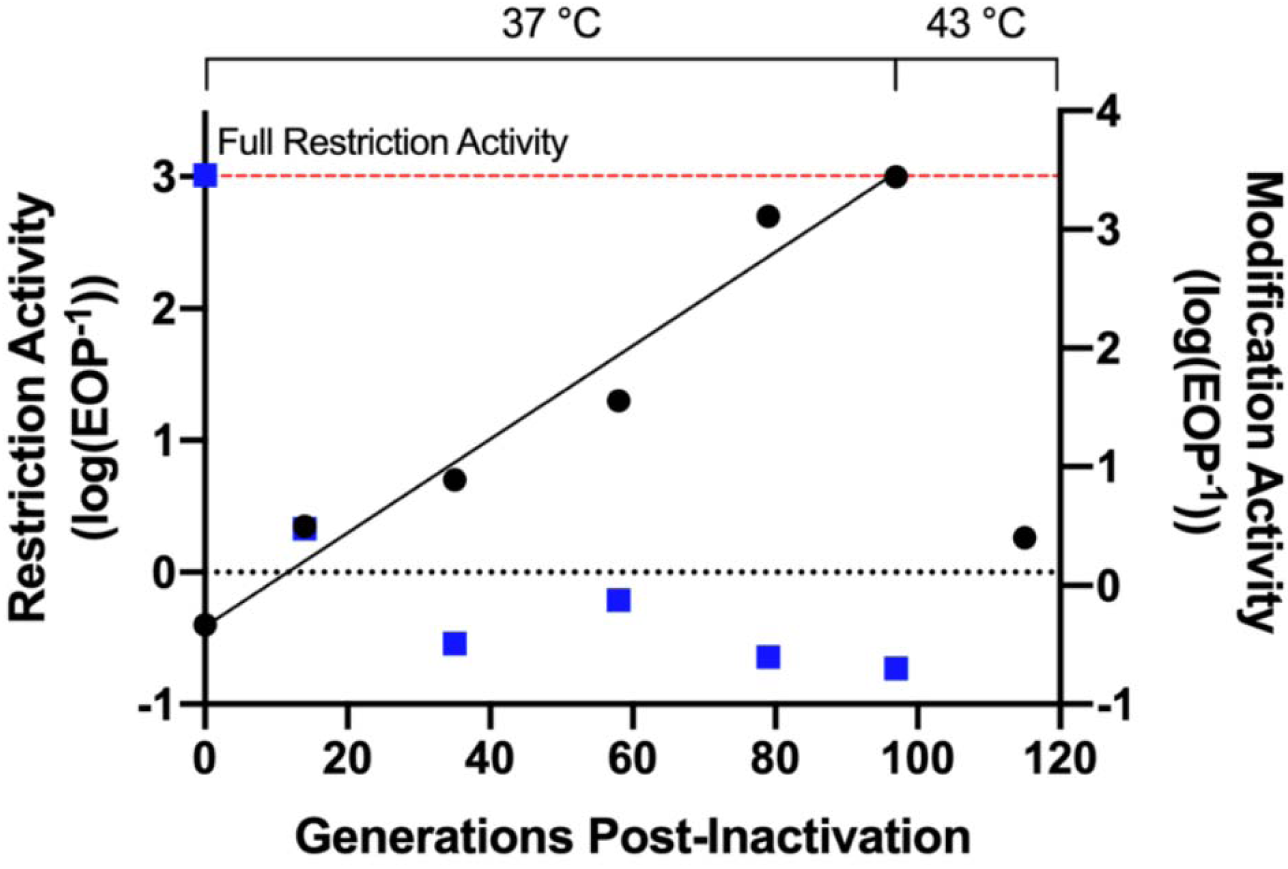
Restriction activity following ON growth at 43 °C, 100 generations recovery at 37 °C, and re-introducing ON growth at 43 °C.

**Figure S4:**
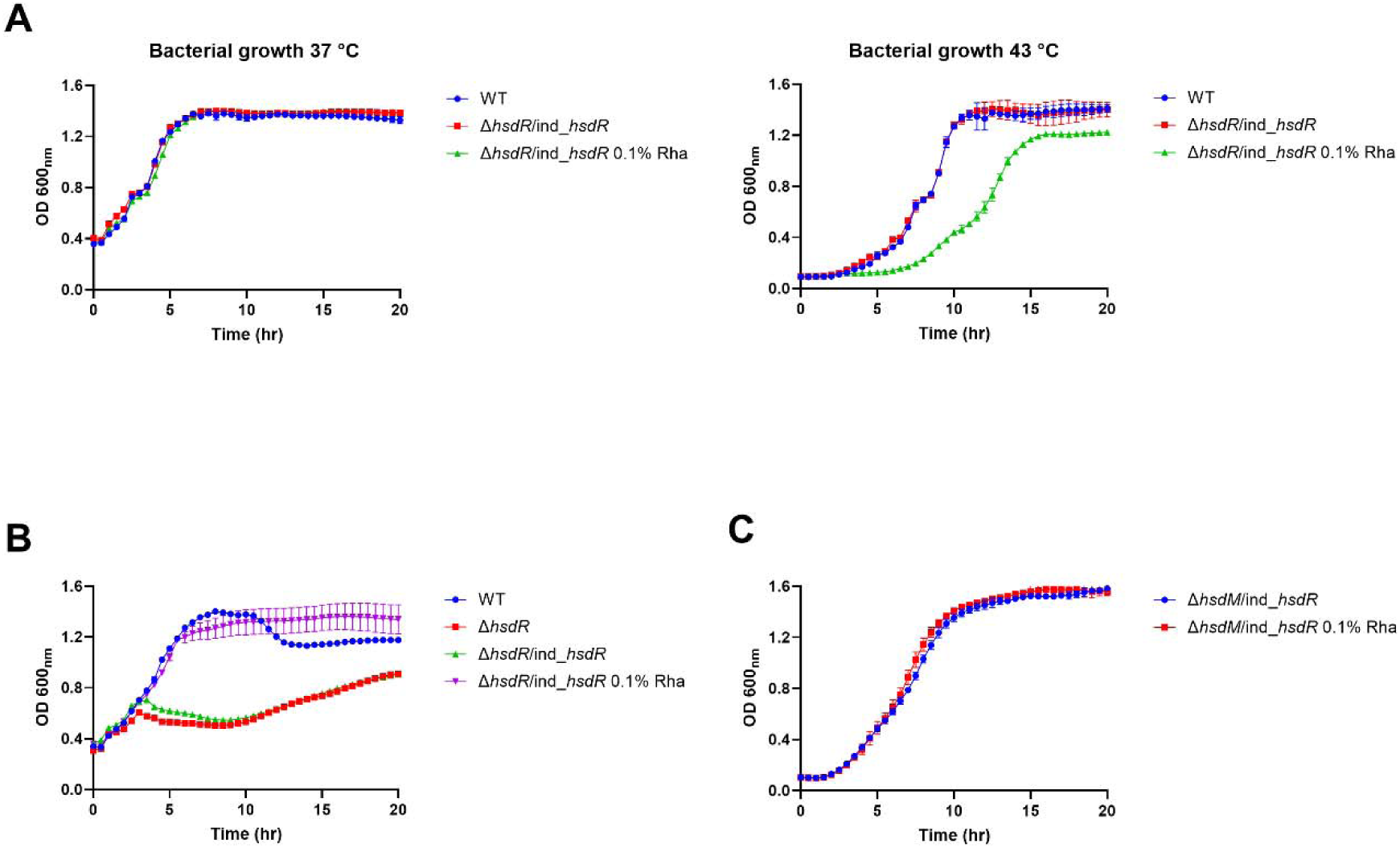
HsdR expression is toxic only at iREN state. **(A)** Growth curve at 37°C following or ON growth at 37 °C (left) or 43°C (right) of WT and strains harboring an inducible copy of *hsdR*, treated with Rha inducer. **(B)** Restriction activity of WT, *hsdR* mutant, and strains harboring an inducible copy of *hsdR* at 37 °C, treated with Rha inducer **(C)** Growth curve at 37°C following ON growth at 43°C (right) of *hsdM* mutant strain harboring an inducible copy of *hsdR* with or without Rha inducer addition. For all the graphs above, inducer was added at t=0.

**Table S1:**
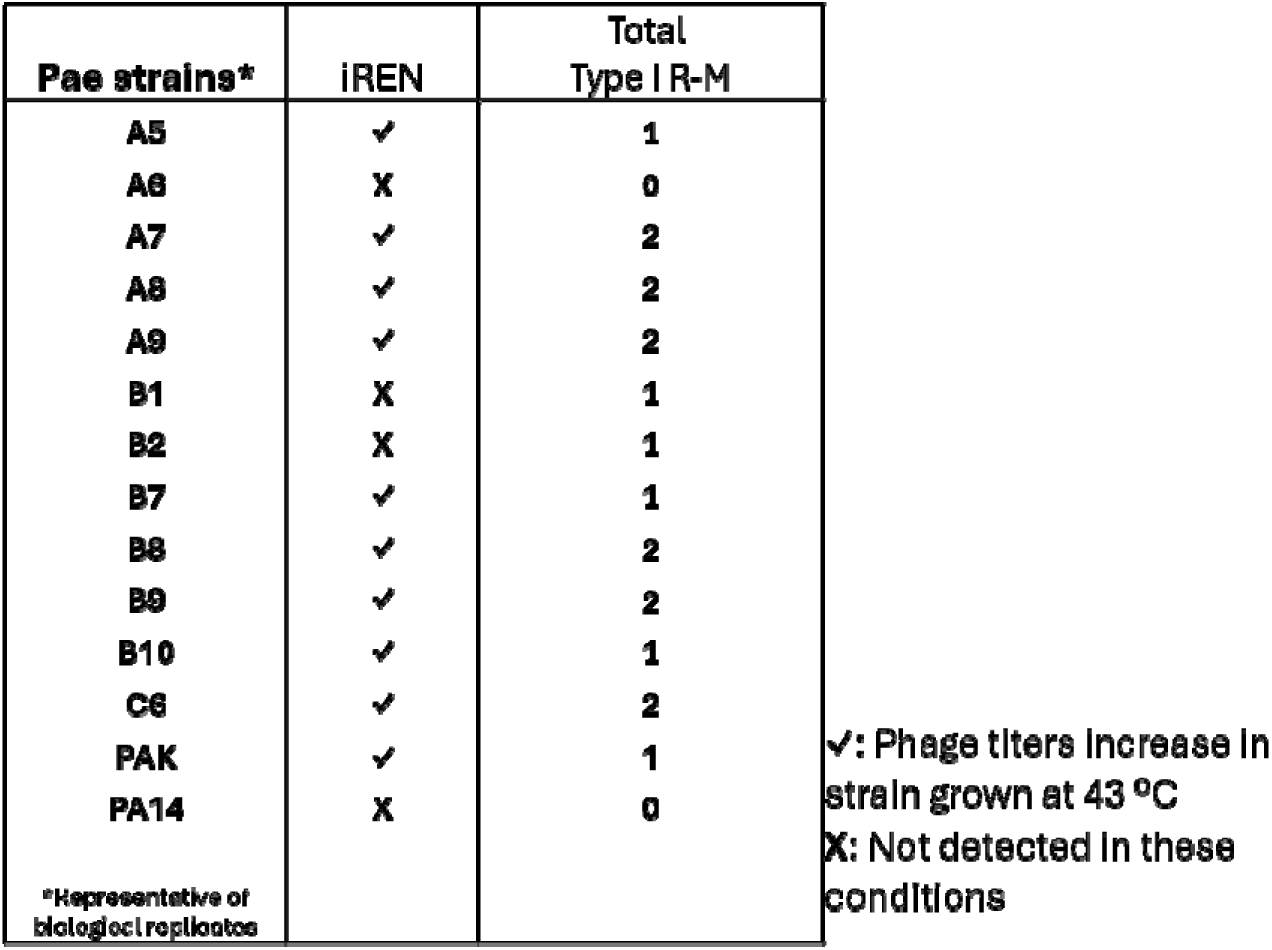
iREN examination in different *P. aeruginosa* strains following growth at 43 °C with indicated genomic type I R-M systems.

**Figure S5:**
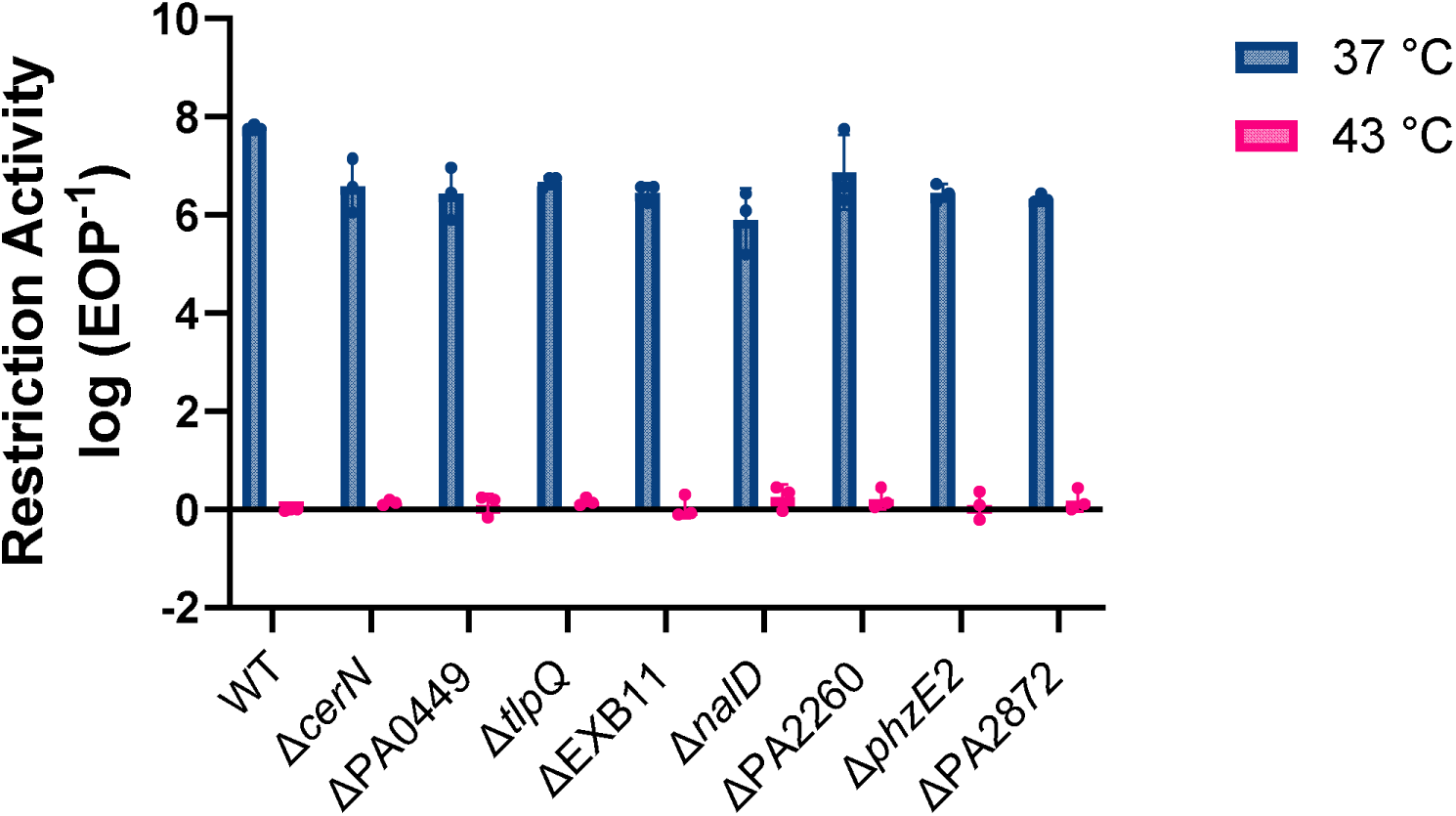
Modification hot spots at 43 °C do not specifically affect iREN. Restriction activity following ON growth at 37 °C (blue) or 43 °C (pink) of WT strain and mutants. Mutants are strains lucking the complete ORF containing the modification site. The graph is the average of three independent biological replicates, with error bars indicating standard deviation.

**Figure S6:**
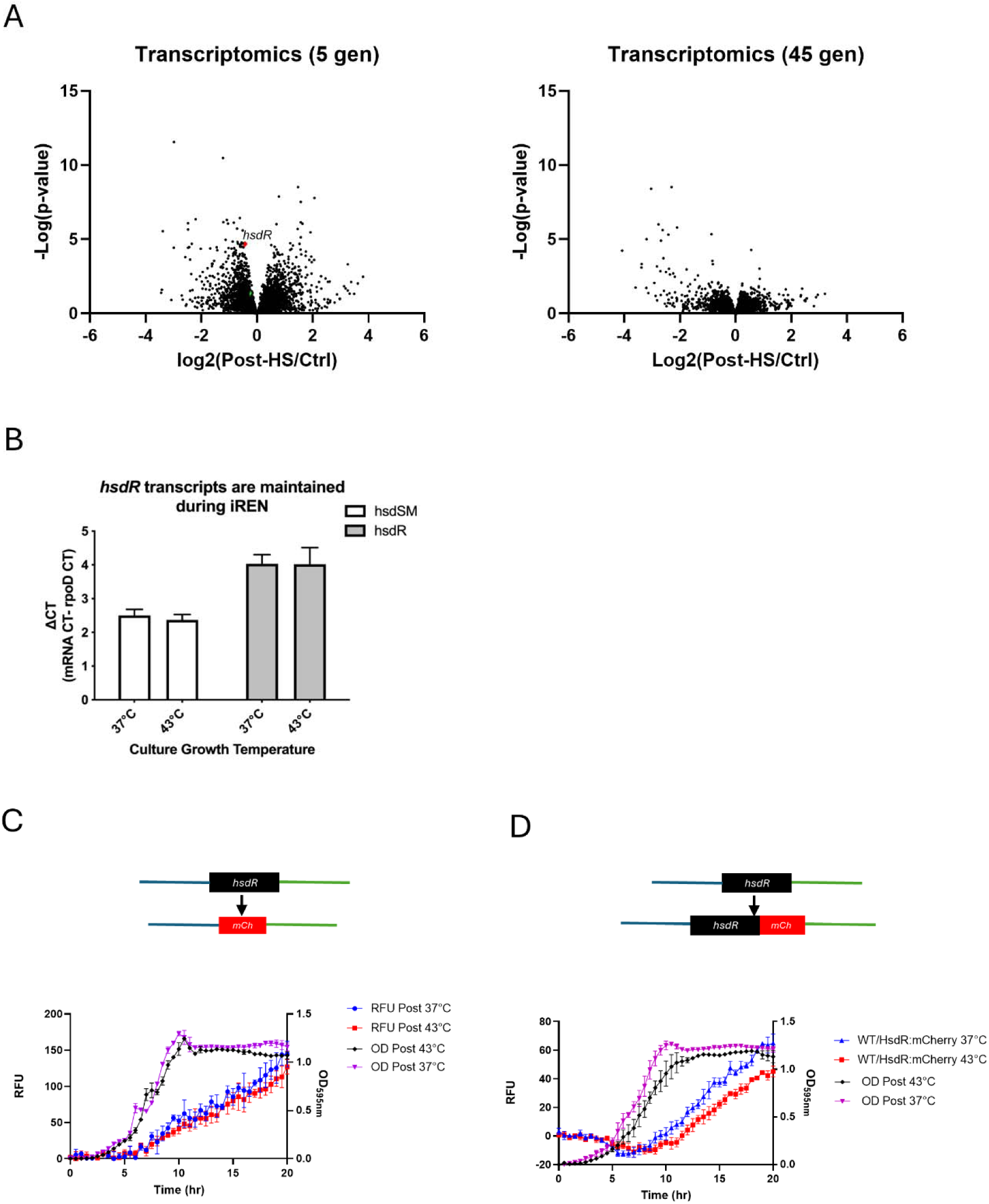
Restriction endonuclease is not transcriptionally regulated. **(A)** Global transcriptomics analysis: Samples taken 5 generations following 43 °C growth and at the end of the recovery phase (45 generations) were compared to normal growth conditions. Marked dots indicate Hsd proteins: HsdR (red) and HsdMS (green). Transcriptional and Translational Reporter Analysis. **(B)** Real-time PCR analysis of *hsd* transcripts following growth at either 37 °C or 43 °C. expression levels are normalized to *rpoD* as a ctrl. **(C)** Fluorescence and absorbance measurements over time of a strain harboring the *hsdR* ORF replaced by *mCherry*, representing transcriptional reporter analysis. **(D)** Fluorescence and optical density measurements over time of a strain harboring *hsdR* fused to *mCherry*, representing translational reporter analysis. The (C) and (D) graphs are the average of three independent biological replicates, with error bars indicating standard deviation.

**Figure S7:**
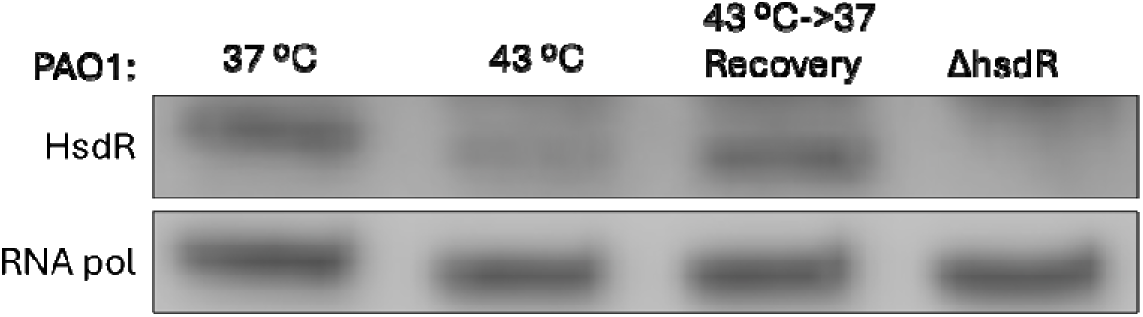
HsdR-specific custom antibody which showed a similar reduction in HsdR levels. Endogenous measurement of Type I R-M proteins, HsdR from WT cells at 37 °C, 43 °C, or recovery at 37 °C for 10 generations by Western blot analysis using custom polyclonal antibodies, RNA pol was used as a loading ctrl.

**Figure S8:**
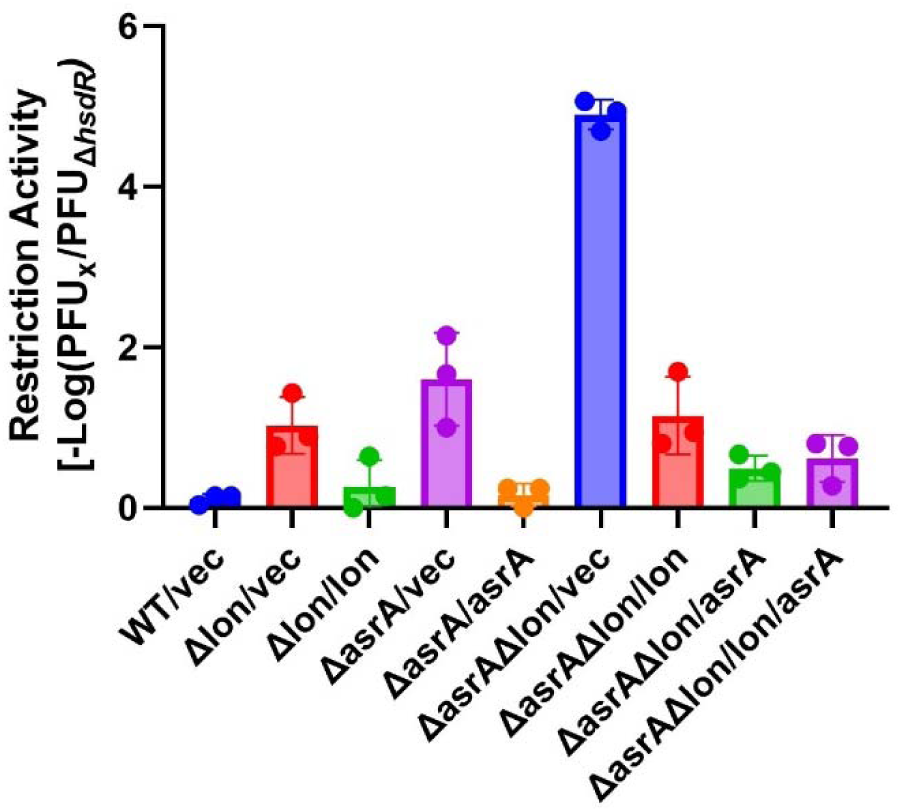
Proteases complementation and HsdR protein levels effect. Protease complementation restores iREN: WT and mutant strains containing either empty vectors or plasmid-expressed protease were examined for restriction activity following ON growth at 43 °C. each dot represents a single biological replicate, with error bars indicating standard deviation.

**Figure S9:**
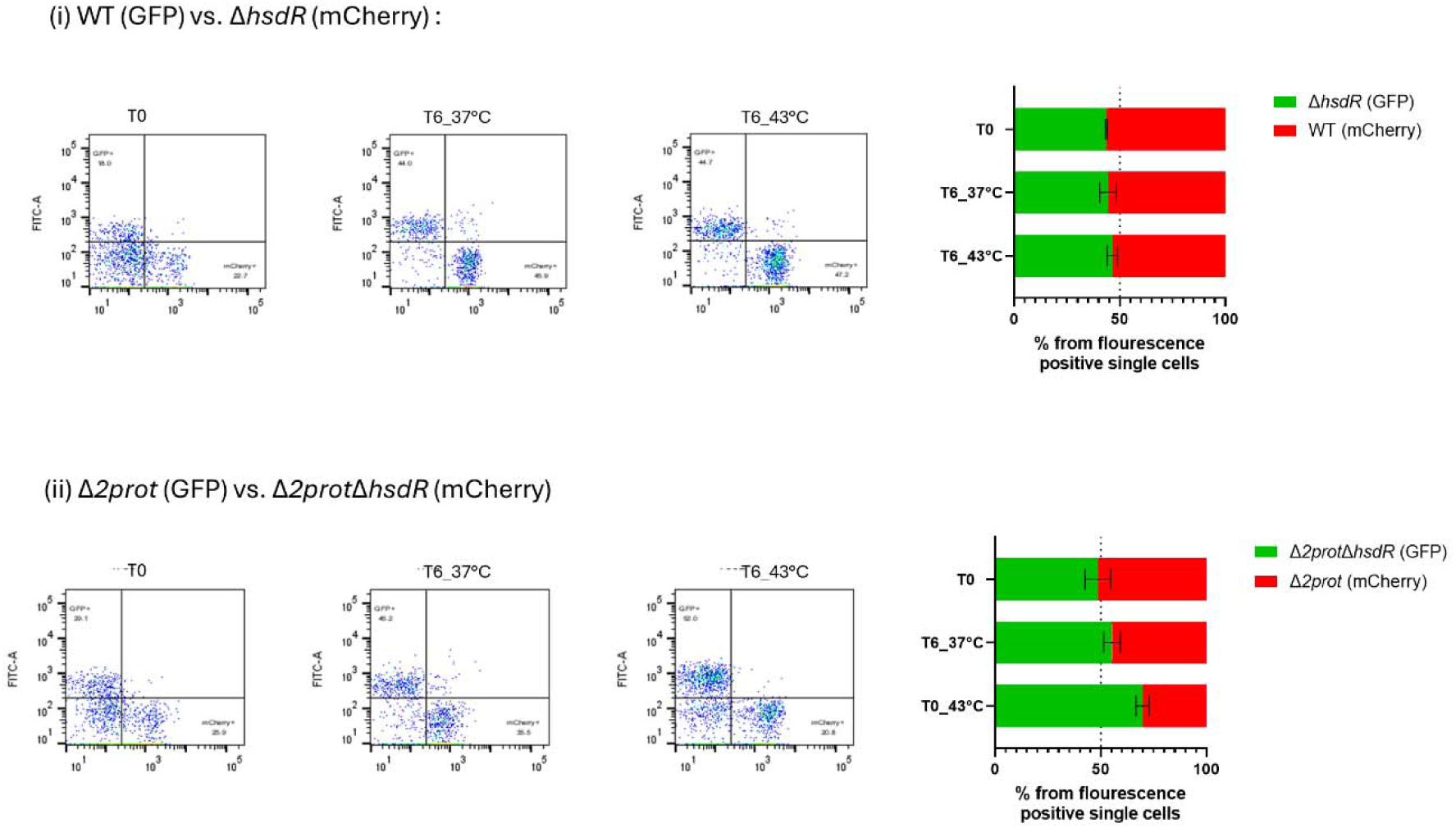
Growth rate and fitness cost for restriction activation in the proteases mutant. **(A)** Growth curves of WT, single mutants, and double 2prot mutants (blue) with additional deletion of *hsdR* (red) at 43°C (up) and following sub-culture to 37°C (bottom). Graphs represent the average of three biological replicates, with error bars indicating standard deviation. **(B)** *hsdR* deletion competition under different conditions, reverse tagging: Flow-cytometry analysis of population variation in mixed WT and Δ*hsdR* strains (top), or double-proteases mutant and double mutant lacking *hsdR* (bottom). Mixtures were grown for six hours before analysis. Representative histograms of the reverse tagging analysis are displayed (left), and population percentages are shown as an average of three biological replicates (right).

**Table S2:**
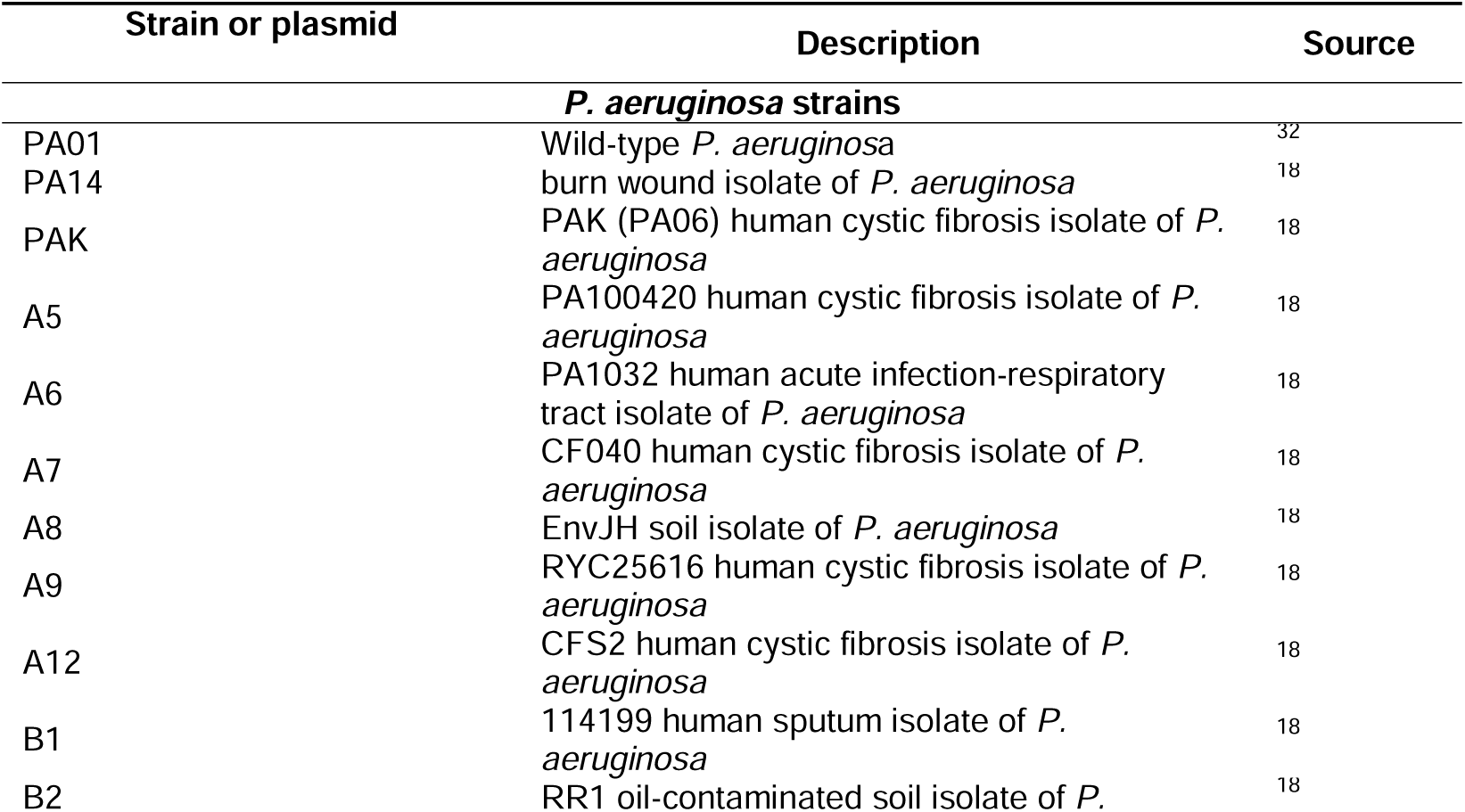
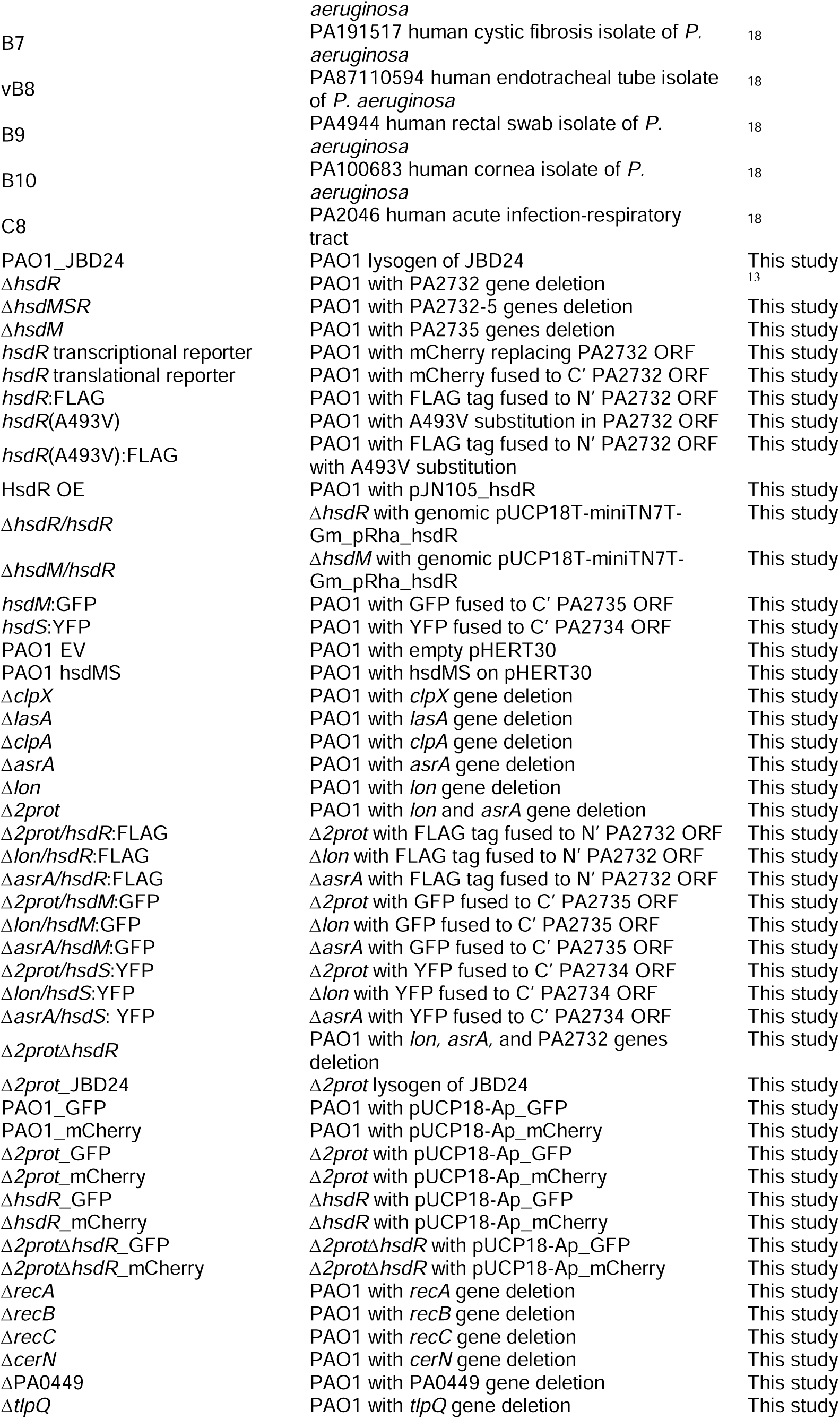
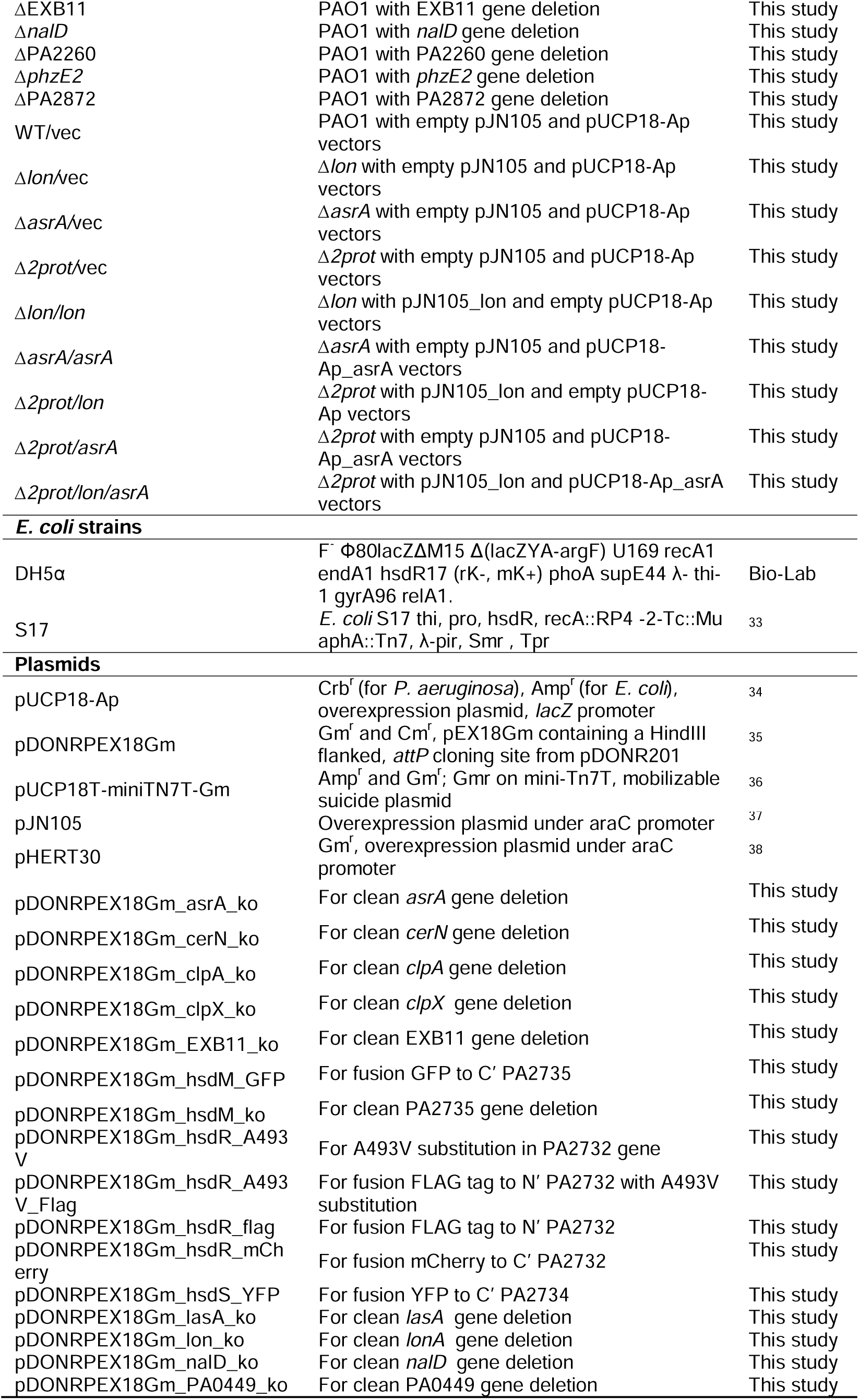
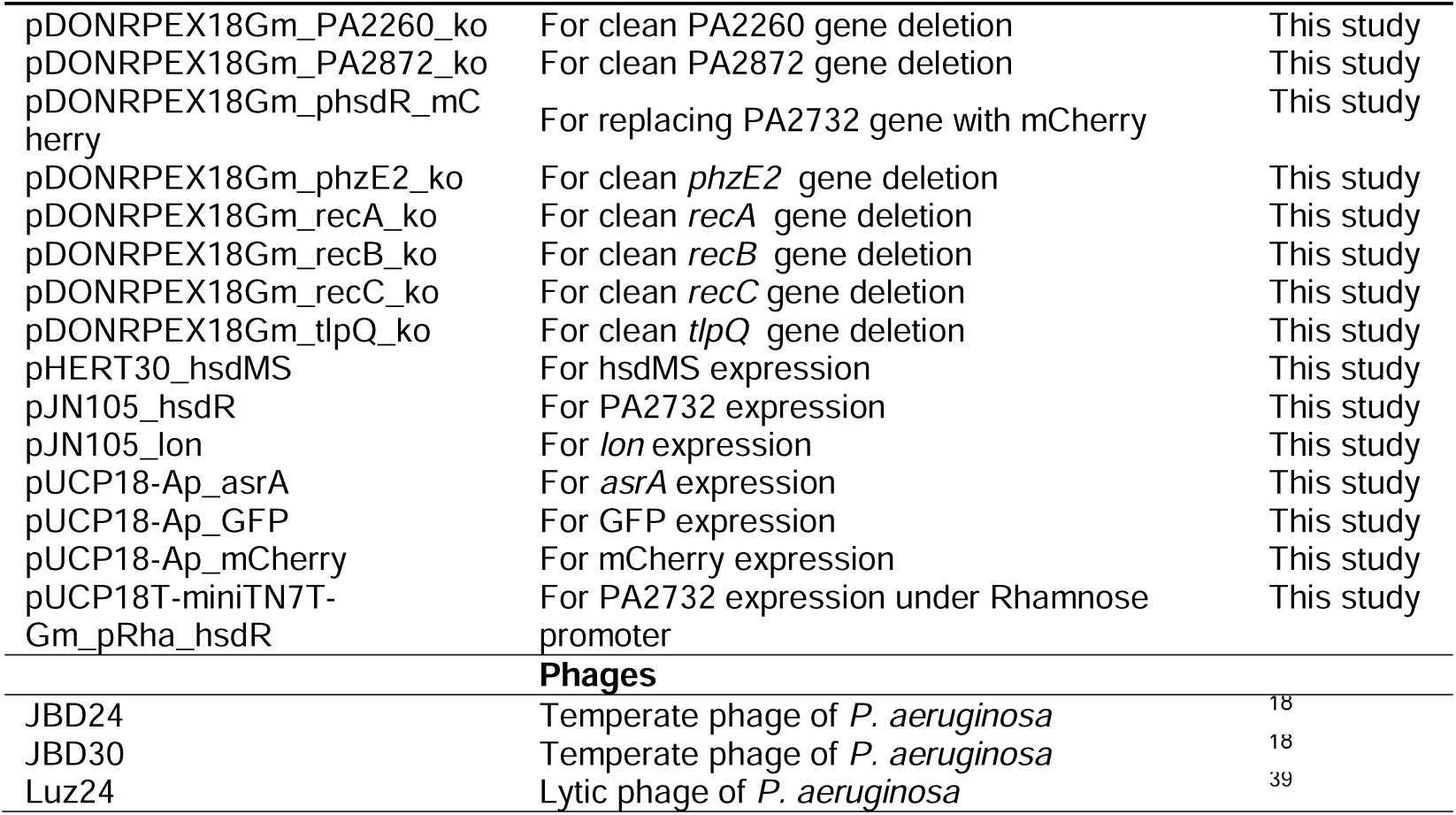
Strains and plasmids used in the study.

